# reComBat: batch-effect removal in large-scale multi-source gene-expression data integration

**DOI:** 10.1101/2021.11.22.469488

**Authors:** Michael F. Adamer, Sarah C. Brüningk, Alejandro Tejada-Arranz, Fabienne Estermann, Marek Basler, Karsten Borgwardt

**Affiliations:** Machine Learning & Computational Biology Lab, Department of Biosystems Science and Engineering, ETH Zurich, Basel, Switzerland; Swiss Institute for Bioinformatics (SIB), Lausanne, Switzerland; Biozentrum, University of Basel, Basel, Switzerland

## Abstract

**Motivation:** With the steadily increasing abundance of omics data produced all over the world under vastly different experimental conditions residing in public databases, a crucial step in many data-driven bioinformatics applications is that of data integration. The challenge of batch-effect removal for entire databases lies in the large number of batches and biological variation which can result in design matrix singularity. This problem can currently not be solved satisfactorily by any common batch-correction algorithm.

**Results:** We present *reComBat*, a regularized version of the empirical Bayes method to overcome this limitation and benchmark it against popular approaches for the harmonization of public gene expression data (both microarray and bulkRNAsq) of the human opportunistic pathogen *Pseudomonas aeruginosa*. Batch-effects are successfully mitigated while biologically meaningful gene expression variation is retained. *reComBat* fills the gap in batch-correction approaches applicable to large-scale, public omics databases and opens up new avenues for data-driven analysis of complex biological processes beyond the scope of a single study.

**Availability:** The code is available at https://github.com/BorgwardtLab/reComBat, all data and evaluation code can be found at https://github.com/BorgwardtLab/batchCorrectionPublicData

## 1 Introduction

Data-driven computational biology greatly depends on the availability of large, integrated data-sets to provide the necessary variety and statistical power for state-of-the-art (SOTA) machine and deep learning, as recently demonstrated by Alpha-Fold (Jumper *et al*., 2021). In particular, an indepth understanding of general trends in expression and transcription profiles are key for important research questions such as overcoming microbial antibiotic resistance, (Gil-Gil *et al*., 2021; Andersson *et al*., 2020) or cancer therapy failure (Kourou *et al*., 2021; Malod-Dognin *et al*., 2019). By mining large databases across studies, it may be possible to identify novel biological mechanisms that cannot be found by studying individual, small-scale experiments alone. This poses a problem shift towards the need for integrating diverse data obtained from numerous independent experiments.

Public databases such as the Gene Expression Omnibus (GEO) (Barrett *et al*., 2013; Edgar *et al*., 2002), include independent studies collected over a large time span, under different biological and technical conditions. Hence, strong batch-effects (i.e. unwanted and biologically irrelevant variation) preclude a comprehensive analysis of pooled data and first need to be mitigated while desired biological variation (referred to in this paper as “(experimental) design”) needs be retained.

Although a range of batch-correction algorithms has previously been suggested (Tran *et al*., 2020; Lazar *et al*., 2012; Rong *et al*., 2020; Chazarra-Gil *et al*., 2021), only a small subset of these remains applicable for this large-scale setting. In particular, most previous algorithms cannot incorporate high-dimensional experimental design information. Our goal for this study is to provide the community with a simple, yet effective extension of the popular and computationally efficient empirical Bayes method (Johnson *et al*., 2006) (ComBat) to account for a large amount of highly correlated biological covariates. ComBat is based on ordinary linear regression and, therefore, will fail if the system is underdetermined.

We benchmark our method on simulated data and provide a real-world application in microarray and bulk RNAsq data, evaluating the impact of culture conditions on the gene expression profiles of *Pseudomonas aeruginosa* (PA). PA is a Gram-negative bacterium with a large genome (Stover *et al*., 2000) that thrives in a variety of environments and has been declared a critical priority pathogen for the development of new antimicrobial treatments (Tacconelli *et al*., 2018). A large range of studies have previously investigated the impact of culture conditions on the gene expression profiles of PA. A comprehensive review of the perturbations caused by the microenvironmental cues is missing as a consequence of the lack of harmonized data allowing for a direct comparison.

The paper is organized as follows. After reviewing relevant literature in Section 1.1 we introduce our *reComBat* algorithm (contribution i) in Section 2 as an extension of the ComBat algorithm to handle highly correlated covariates. In the second part of Section 2 we address the issue of assessing the efficacy of the batch-correction by introducing a large variety of evaluation metrics (contribution ii). In Section 3 we benchmark *reComBat* against a selection of SOTA batch-correction methods on simulated and real-world data. Finally, we present a large, harmonized data-set of PA expression profiles in response to different microenvironmental cues (contribution iii). We conclude Section 3 by demonstrating, as a proof of concept, the biological validity of the harmonized data-set. Section 4 comprises of a discussion and outlook.

### 1.1 Related Work

A variety of batch-correction methods has previously been suggested for bulk and single cell sequencing data (see e.g. (Lazar *et al*., 2012; Tran *et al*., 2020; Yu *et al*., 2021)). Here, we focus on batch-correction of bulk data which can generally be divided into the following categories:

#### Normalization to reference genes or samples

Algorithms, such as cross-platform normalization (Shabalin *et al*., 2008) or reference scaling (Kim *et al*., 2007), which employ references, are infeasible in the public data domain: “reference” or “house keeping” genes don’t exist for some organisms, particularly microbes, eliminating these as common ground for batch-effect correction. Given a large public data-set, overlapping samples or common reference experiments are unlikely.

#### Discretization methods

Approaches that discretize expression data into categories (e.g., “expressed” vs. “not expressed”) can be hard to implement rigorously without a relevant control. Furthermore, the information loss due to discretization may affect the results of any advanced downstream analysis of the harmonized data.

#### Location-scale adjustments

These methods adjust the mean and/or variance of the genes, e.g by standardization (Li and Wong, 2001) or batch mean-centering (Sims *et al*., 2008). This only works if the batch-effect is a simple mean/variance shift and does not account for additional confounders. One of the most popular location-scale method is the empirical Bayes algorithm, ComBat (Johnson *et al*., 2006). Despite reasonable success for the correction of local, i.e. within one experiment, or moderate (i.e. comprising few, biologically correlated) batch-effects most location-scale adjustment methods either provide insufficient correction in the presence of strong batch-effects (e.g. standardization) or are unable to account for highly correlated design features (e.g. ComBat).

#### Matrix factorization

This approach builds on decomposition such as principal component analysis (PCA) or singular value decomposition (SVD) (Alter *et al*., 2000) to identify and remove factors characterizing the batch. While this can work in small scale experiments, it is unclear how to apply these methods when there is strong confounding of batch and biological variation. A tangential approach to matrix factorization is to estimate unwanted variation via surrogate variables (SVA) (Lazar *et al*., 2012). Since in our setting we assume that we know all sources of variation, we do not consider SVA.

#### Deep learning based

Recently, nonlinear models, often based on neural/variational autoencoders or generative adversarial networks (GAN), have gained popularity (e.g. normAE (Rong *et al*., 2020), AD-AE (Dincer *et al*., 2020), scGen (Lotfollahi *et al*., 2019), (Ghahramani *et al*., 2018)). This class of models aims to find a batch-effect-free latent space representation of the data e.g. via adversarial training. While an advantage of these methods is their flexibility to account for batches, but also desired biological variation, a major drawback may be that the batch-effect is only removed in a low-dimensional latent space. Downstream analysis is necessarily constrained (Dincer *et al*., 2020; Rong *et al*., 2020). scGen is a notable exception as it provides a direct normalization at gene expression level. However, large data-sets are required and, in the absence of ground truth, the risk of overcorrection should be considered in addition to increased computational complexity.

## 2 Approach

In this section we introduce the mathematical tools and start by defining our modification to the popular ComBat algorithm, *reComBat*, before introducing a range of possible evaluation metrics to gauge the efficacy of data harmonization.

### 2.1 Classical: ComBat

ComBat (Johnson *et al*., 2006) is a well-established batch-correction algorithm employing a three-step process.

1. The gene expressions are estimated via an ordinary linear regression and the data is standardized.
2. The adjustment parameters are found by empirical Bayes estimates of parametric or non-parametric priors.
3. The standardized data is adjusted to remove the batch-effect.

The ComBat algorithm has seen many refinements and applications (see e.g. (Čuklina *et al*., 2021; Müller *et al*., 2016; Zhang *et al*., 2020)). However, most data-sets have still been small and did not come with an extensive design matrix. When the design matrix becomes large (many covariates) and sparse, unexpected issues can arise in step 1 of the algorithm. To illustrate the classic algorithm, we use the slightly modified ansatz of (Wachinger *et al*., 2021),

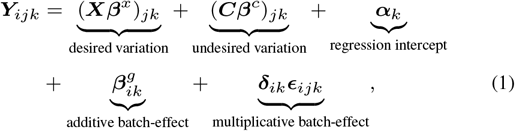

where ***Y***_*ijk*_ is the gene expression of the *k*^th^ gene in the *j*^th^ sample of the *i*^th^ batch. The matrices ***X*** and ***C*** are design matrices of desired and undesired variation with their corresponding matrices of regression coefficients ***β***^*x*^ and ***β***^*c*^. ***α*** is a matrix of intercepts, and ***β***^*g*^ and ***δ*** parameterize the *additive* and *multiplicative* batch-effects. The tensor ***ϵ*** is a three-dimensional tensor of standard Gaussian random variables. Note, that we implicitly encode batch-and sample-dependency by dropping the relevant indices, i.e. ***β***^*g*^ depends on the batch and gene, but is constant for each sample within the batch.

In the first step of the algorithm the parameters ***β***^*x*^, ***β***^*c*^, and ***α*** are fitted via an ordinary linear regression on

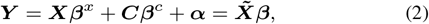

Where 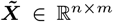, where *m* is the number of features and *n* is the number of samples. Note, that this formulation is equivalent to redefining ***Y*** ∈ ℝ ^*n*×*g*^, where *g* is the number of genes, and subsuming the batch and ***C*** features into 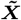. The intercept ***α*** is inferred via the relation 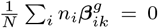 (Johnson *et al*., 2006), where *n*_*i*_ is the number of samples in batch *i*, ***β***_***ik***_ is the regression coefficient of batch *i* for gene *k* and *N* is the total number of samples. For ease of notation, in the remainder of this paper we will use this equivalent formulation.

Once, the model is fitted, the data is standardized, then the batch-effect parameters, 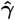 and 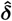 are estimated using a parametric or non-parametric empirical Bayes method. Finally, the data is adjusted. For details, please refer to the original publication (Johnson *et al*., 2006).

### 2.2 Novel contribution: *reComBat*

#### Problem statement

Using standard results for ordinary linear regression, we know that if the matrix 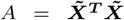 is positive-definite, the optimization of (2) is strictly convex. However, if ***A*** is singular a unique-solution the the regression does not exist. Hence, if ***A*** is rank-deficient, i.e. the system is underdetermined, ComBat will not necessarily arrive at a unique solution. Our goal in this work is to provide a computationally efficient solution for this problem to make the empirical Bayes method applicable also to large-scale public data harmonization.

Given the popularity of ComBat this issue does not seem to be encountered frequently. One possible explanation is that the sources of biological variation that are usually considered within the same experiment are limited and well-chosen. When integrating entire databases, however, the sources of biological variation are manifold and these can often only be encoded as categorical variables. One prominent example is considering all uploaded experimental data of a particular pathogen, which can result in hundreds of unique experimental conditions, some potentially highly correlated with other metadata. Encoding these as one-hot categorical variables creates a sparse, high-dimensional feature vector and, when many such categorical features are considered, then *m* ≈ *n*. If, either *m* > *n*, or strong batch-design correlations exist, then, even for large-scale integration, ***A*** may be rank deficient.

To mitigate this issue, we propose a modification of the estimation of gene expression profiles by a linear model (step 1 of the ComBat algorithm) by fitting the elastic net model -a standard approach from linear regression theory

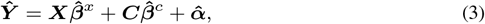

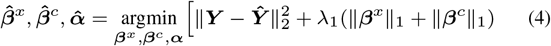

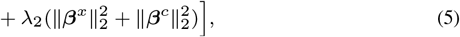

where ∥·∥_*p*_ denotes the *𝓁*_*p*_ norm, and λ_1_ and λ_2_ are the LASSO and ridge regularization penalties. Due to this regularizing modification of the algorithm we call our approach **re**gularized-ComBat, in short *reComBat*. Both, parameter fitting using the Empirical Bayes methods, and parameter adjustment on the standardized data follow the above outline for the ComBat algorithm. Note that *reComBat* essentially replaces a linear regression with a regularized regression and, hence, the increase of computational complexity of *reComBat* over ComBat is negligible.

The *reComBat* algorithm can be summarized in the following pseudo-code.

##### Algorithm 1

reComBat

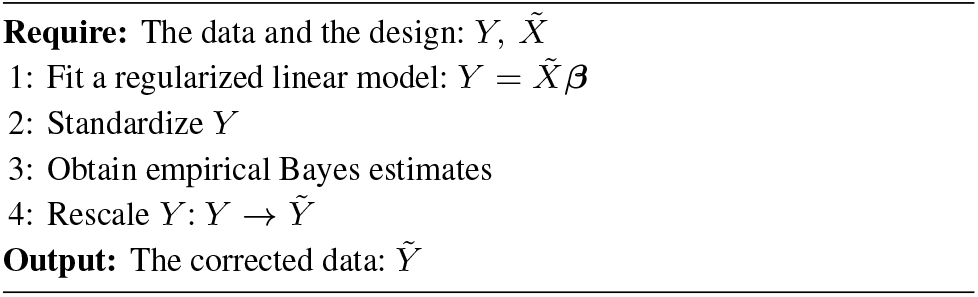

### 2.3 Evaluation metrics

A detailed description and definition of all evaluation metrics employed to score batch correction efficacy is provided in supplement A. We included classifier-based (logistic regression-based balanced accuracy and F1-score, Linear Discriminant Analysis (LDA) score), cluster-based (minimum separation number, cluster purity, Gini impurity), and sample distance-based (Distance Ratio Score (DRS), Shannon entropy) metrics.

## 3 Experiments

In this section, we apply *reComBat* to simulated and real-world microarray and bulkRNAsq data. We show quantitatively and qualitatively that *reComBat* is successful in removing substantial batch-effects while retaining biologically meaningful signal.

### 3.1 Experimental data

A detailed description is given in supplement B. We first evaluate the approaches on synthetic data with singular design matrix and test a range of hyperparameter combinations for data generation (number of samples (100-2000), batches (3-100), design matrix features (3-20), relative disturbance size of metadata to batch (0.01-20), number of Zero-Hops (5-40)) and score run time, LDA score, Shannon entropy and cluster purity as a function thereof w.r.t. the ground truth. Additionally, data for 887 (114 batches, 39 Zero-Hops, see Table S1) microarray and 340 bulkRNAsq samples (32 batches, 12 Zero-Hops, see Table S2) was collected from the GEO, SRA and ENA data bases (Barrett *et al*., 2013) with relevant metadata characterizing experimental design (culture conditions, PA strain). The obtained microarray design matrix is singular, whereas the RNA design matrix is not-singular, however, ill-conditioned.

### 3.2 Batch-correction methods

We tested our approach against a representative sample of baseline methods, in particular, standardization, marker gene elimination, principal component elimination, ComBat, Harmony (Korsunsky *et al*., 2019) and scGen. Details on these methods can be found in the supplement C.

For *reComBat*, we used parametric priors for the empirical Bayes optimization and tested a variety of parameters including pure LASSO (λ_2_ = 0), pure ridge (λ_1_ = 0), and the full elastic net regression. The range of regularization strengths tested were all possible combinations (except for (0, 0)) of λ_1_ ∈ {0, 10^−2^, 10^−1^, 1} and λ_2_ ∈ {0, 10^−10^, …, 10^−1^, 1}. Note that smaller values of λ_1_ yielded numerical instabilities.

### 3.3 Hyperparameter optimization results

A hyperparameter screen to optimize regularization strength and type on the default simulated, microarray and bulkRNAsq data yielded best results when ridge regression was used (λ_1_ = 0) with λ_2_ ≤ 0.001 (see supplement D). The specific regularization parameter only had a minor influence and we continued with λ_2_ = 10^−9^. We observe that stronger, particularly LASSO, regularization achieves superior batch heterogeneity at the cost of decrease in Zero-Hop uniformity in real-world data. Notice that LASSO-*reComBat* performs implicit feature selection due the *𝓁*_1_ regularization. This could hint to the fact that more balanced feature weighting (as provided by ridge-*reComBat*) is beneficial. In the following we present results only for ridge *reComBat*.

### 3.4 Evaluation on synthetic data

We benchmark *reComBat* on simulated data against popular batch-correction methods. Figure 1 A,B shows the simulated ground truth distribution together with the distribution after applying batch-effects, and following data harmonization with *reComBat*. The ground truth results in terms of Zero-Hop clusters were qualitatively well reproduced by *reComBat*. Quantitative results in terms of LDA score difference to ground truth (see supplement E for Shannon entropy, Gini impurity and cluster purity) are shown in Figure 2A as a function of different data generation hyperparameters for the investigated correction methods. We observe that *reComBat* and scGen outperform Harmony and simple correction (PC or marker gene elimination, standardization). Notably, if scGen is trained with Zero-Hop labels its performance is greatly improved, however, also prone to overfitting (positive LDA score differences). We only observe degradation of *reComBat* performance for smaller data-sets of 100 samples (given 10 Zero-Hops). Run time was generally very quick and favorable for *reComBat* compared to Harmony, or scGen (trained on GPU).

**Fig. 1.**
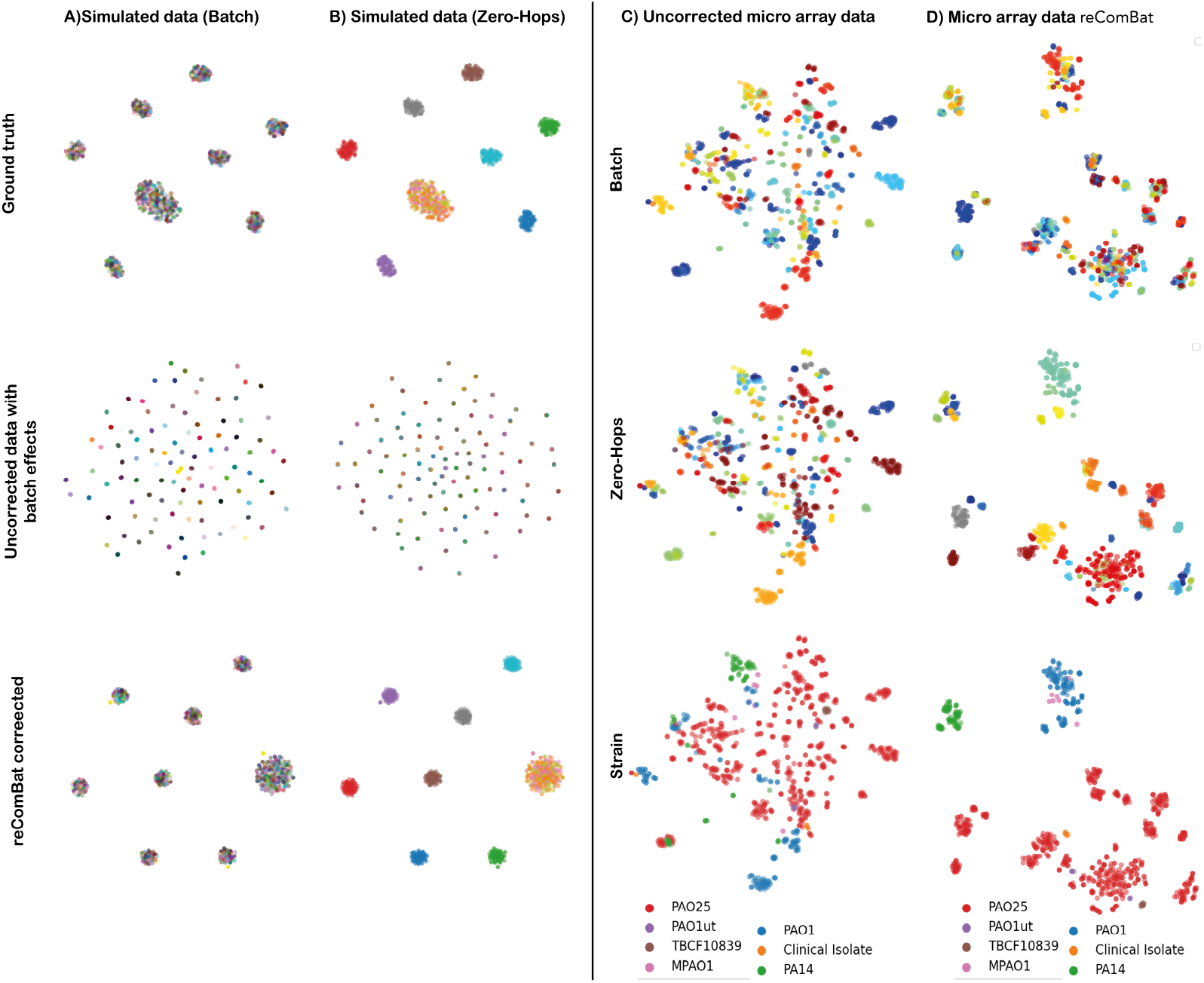
t-SNE plots of the simulated (A, B) and microarray (C, D) data-sets. For simulated data we show ground truth (top), uncorrected (middle) and reComBat (λ_1_ = 0, λ_2_ = 10^−9^) corrected (bottom) results. (Un)Corrected microarray data are colored by batches (top), Zero-Hops (middle), and microbial strain (bottom). Color scales do not reflect proximity of the relevant batches or Zero-Hops.

**Fig. 2.**
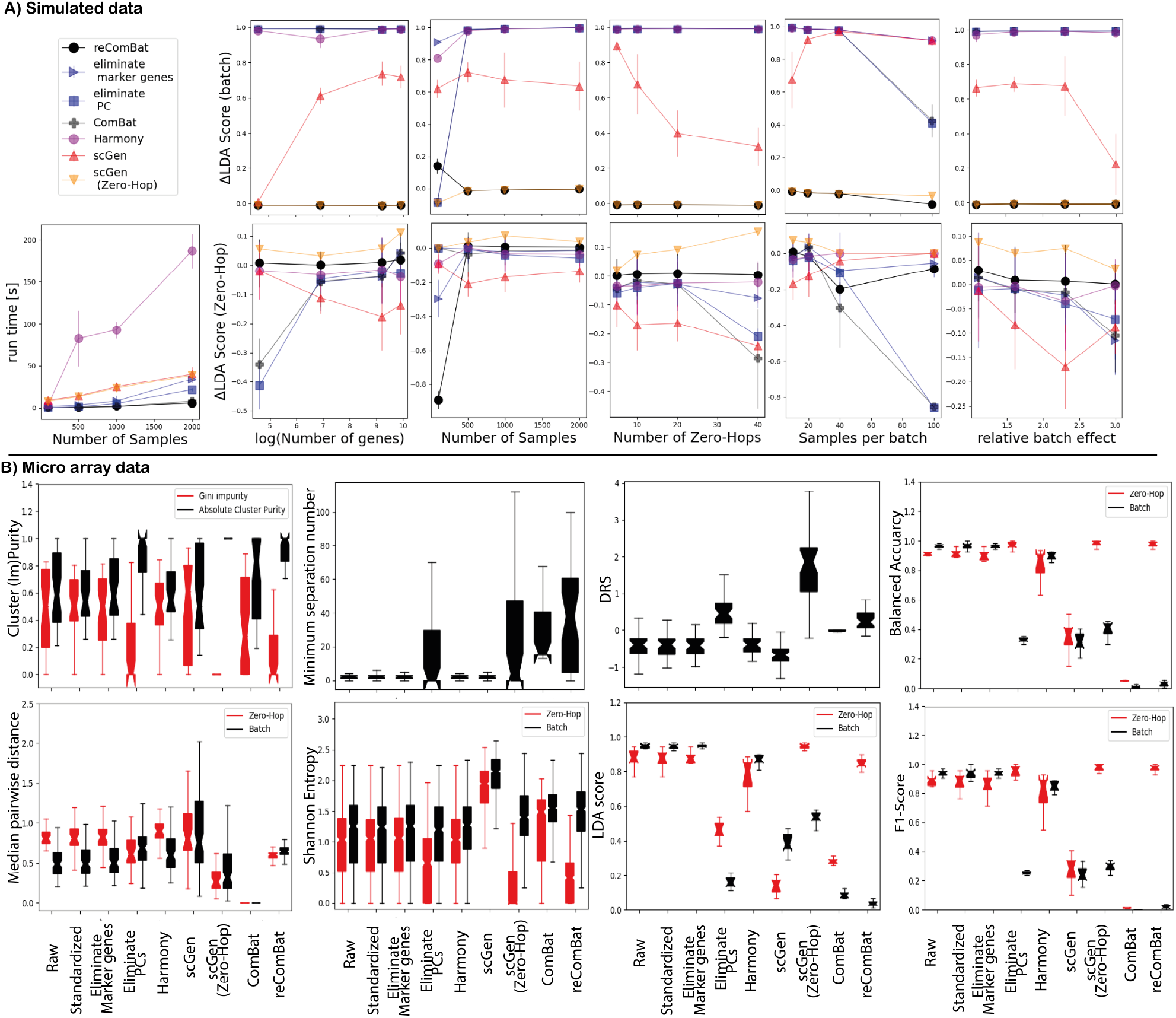
A) Overview over results based on different simulated data-sets scored in terms of LDA score difference to ground truth for batch and Zero-Hops. Results represent mean values and standard deviations over 10 independent repeats. B) Evaluation metrics scoring the impact of batch-effects by evaluating the variety of different batches and/or Zero-Hops of the (un-) corrected microarray dataset. Box plots represent the lower and upper quartiles (box) together with the median (central dents) and full range (whiskers) over all samples, clusters, or Zero-Hops depending on the relevant metric. LDA scores and LR classification performance are reported over ten cross validation folds.

### 3.5 Experimental benchmarking of *reComBat*

We show quantitatively and qualitatively that *reComBat* is successful in removing substantial batch-effects while retaining biologically meaningful signal in real-world data, too. Figure 1 C,D gives an overview of the uncorrected and *reComBat* corrected microarray data colored by batch, Zero-Hops, and microbial strain. Uncorrected data clusters by batch, indicating the presence of batch-effects, whereas clustering by biologically meaningful variation (e.g. by strain or Zero-Hop) is observed after correction. Additional overviews of t-SNE embeddings of batch-corrected expression data for all baseline models and data, colored by all design matrix elements are provided in supplement F.

We compared our baselines to the best performing *reComBat* model based on all evaluation metrics (supplement C) in Figure 2B. In terms of gauging the metrics themselves for the ability to detect batch-effects, we conclude that classifier-based metrics provide the clearest overview. Shannon entropy can detect a larger spread in batch vs. Zero-Hop entropy, however, the findings may strongly vary by the specific subset. It can also be argued that entropy strongly depends on the choice of the number of nearest neighbors. Likewise, the median pairwise distance and DRS metrics show some ability to detect batch-correction, but due to the strong dependency on the individual Zero-Hop the spread in values may be large. The minimum separation clustering clearly shows when a batch-correction can be considered effective. However, due to repeated clustering, calculation of minimum separation number is computationally far more expensive than distance-based metrics. A good mid-point between classifier-and cluster-based evaluation are cluster-purity measures, which show good resolution and manageable dependency on the Zero-Hop.

Data standardization, and marker gene elimination only had a minor, insignificant (all Mann-Whitney U-Test p-values *>* 0.05) effect when compared to the raw data, independent of the underlying metric and data-set. Despite, markedly different results compared to the uncorrected baseline, Harmony could not achieve sufficient batch-correction characterized by poor performance in classifier and cluster-based metrics throughout. We suggest that the large number of design matrix elements and comparably strong batch-effect could lead to this result. Importantly, *reComBat* achieved good scores throughout all evaluation metrics for all data-sets (bulkRNAsq given in supplement), whereas performance of other correction methods such as PC elimination, scGen, and ComBat varied depending on data and metric. As expected, singularity of the design matrix led to poor performance of ComBat (microarray data), whereas bulkRNAsq data with a non-singular design matrix achieved the best results for this method. For scGen it was key to provide information on Zero-Hops as labels to the algorithm (scGen(Zero-Hop)), whereas simply relying on design matrix covariates led to poor correction.

#### 3.5.1 Characterization of the harmonized microarray data-set

In order to preclude overcorrection (Zindler *et al*., 2020) in the absence of ground truth, we demonstrate that biologically meaningful expression profiles are retained after batch-correction. As representative examples we analyzed data subsets by oxygenation status, culture medium richness, growth phase, or clinical vs. laboratory PA strains in our microarray data-set (supplement G). We identify Zero-Hop marker genes driving the differences between selected pairwise comparisons and assess their relevance to underlying biological pathways. Pathways previously known to be important in the relevant culture conditions were identified. For instance, when comparing standard to hypoxic conditions, we find that genes involved in aerotaxis (Hong *et al*., 2004), Fe-S cluster biogenesis (Romsang *et al*., 2015) and iron acquisition ((Glanville *et al*., 2021; Hannauer *et al*., 2012) are major drivers of differences. When comparing cultures in exponential to stationary phase under hypoxia conditions, genes involved in pyoverdin (Drake *et al*., 2007; Vandenende *et al*., 2004) and pyochelin (Ankenbauer and Quan, 1994; Reimmann *et al*., 2001) biosynthesis and transport, iron starvation (Alontaga *et al*., 2009; Hassett *et al*., 1997; Zhao and Poole, 2000) and quorum sensing (Kim *et al*., 2012) were relevant. Finally, for a comparison between the laboratory strain PAO1 vs. clinical isolates we find cup genes (PA4081-PA4084, PA0994) that are involved in motility and attachment and with this in biofilm formation (Ruer *et al*., 2007). This indicates a difference in attachment between those strains that might be coming form the environment the strains have adapted to grow in (laboratory vs. patient). In all cases, a large amount of hypothetical genes of unknown function also flagged up - an expected observation as roughly two thirds of the genes encoded in the PA genome have an unknown function. The harmonized data-set hence serves for hypothesis generation motivating further (experimental) validation.

## 4 Discussion

Public databases play an increasingly important role for data-driven meta-analysis in computational biology. Despite great efforts to harmonize data collection, considerable, yet unavoidable, biological/technical variation may mask true signal if data are pooled from several sources. To draw generalizable conclusions from agglomerated data, it is essential to correct such batch-effects in a setting where overlapping samples, or standardized controls, are unavailable. When large numbers of (*>* 20) batches coincide with desired biological variation, a range of standard batch-correction algorithms are inapplicable. We would like to stress that this evaluation scenario greatly differs from previously analyzed batch-correction settings where comparably few (2-5) batches with large number of overlapping samples were included, or comparably small batch-effects within a single study were corrected (Tran *et al*., 2020). A key assumption of meta-analysis of published data is the coincidence of “batch” with “study”. Given the substantial manual data curation to extract relevant design matrix information for experimental data the variety of data types (microarray, bulkRNAsq) and organisms (PA) assessed in addition to simulated data was limited. *reComBat* is a simple yet effective, means of mitigating highly correlated experimental conditions through regularization and we compared various elastic net regularization strengths for the purpose of meta analysis based on large-scale public data. We note that given the large number of batch-correction methods available, we only included representative examples for key concepts, including deep, non-linear models (scGen), Harmony, marker gene and PC elimination to benchmark our linear empirical Bayes method.

In case of a singular design matrix *reComBat* outperformed standard approaches, including data standardization, PC and marker gene elimination, Harmony, and scGen if no additional information regarding the evaluation endpoints (here Zero-Hops) was given to either of the methods. We demonstrate not only the superiority of *reComBat* compared to these baselines but, by providing a large variety of evaluation metrics, also give a notion of overall performance.

Importantly, in any large-scale meta-analysis setting, a ground truth is unavailable. Here, biological validation is essential prior to hypothesis generation and we demonstrate this for *reComBat*. Due to this fact we excluded some popular deep models (e.g. normAE(Rong *et al*., 2020), AD-AE (Dincer *et al*., 2020)) from this study as they only provide a latent representation rather than direct correction at gene expression level. These methods would likely provide good batch-correction, however, downstream analysis via e.g. differential gene expression is impossible. There is also growing concern that batch-correction, particularly deep models, may overcorrect and remove biological signal. Although synthetic data addresses this challenge, algorithm performance varies between use-cases and the risk of overcorrection persists. We demonstrate this based on scGen(Zero-Hop) in our benchmark. Both scGen and Harmony (in the published python packages) do not allow for a separation of batch-correction training and validation to test for overfitting by cross-validation -*reComBat* indeed could be used in a cross-validation setting. Notably, in case of e.g.large-scale single cell RNA sequencing, the situation may in fact be favorable for nonlinear approaches -which is not the setting of interest here.

It was possible to show that *reComBat* retained biologically meaningful target pathways identified in a literature-based validation. By mining the harmonized data-set, we can now perform comparisons that have, to the best of our knowledge, never been directly performed before for the purpose of hypothesis generation. For instance, when we compare growth in LB with growth in media that have fewer nutrients, we find that several nutrient (Bains *et al*., 2012; Ball *et al*., 2002; Faure *et al*., 2014; Jones *et al*., 2021; Lewenza *et al*., 2011; Quesada *et al*., 2016) and metal (Alontaga *et al*., 2009; Merriman *et al*., 1995) uptake pathways are deferentially regulated. Experimental validation of the proposed findings is key in confirming information on the underlying biological mechanisms. With *>*5000 citations ComBat is one of the most popular batch-correction methods today applied to a large variety of data types and organisms (Wachinger *et al*., 2021). In this study we showed how an adaptation of this popular algorithm can drastically increase its usability. ComBat benefits from low computational cost, rigorous underlying theory, interpretability, and is easy to apply in practice. We specifically want to recommend *reComBat* in a setting of comparably strong batch-effects and diverse experimental designs as are frequently observed within publicly sourced data from different laboratories. We acknowledge the small methodological differences between ComBat and *reComBat* but stress the importance of this adaptation to make a well-established method suitable for large-scale public data integration. By publishing *reComBat* as a python package^1^ our method is readily available to the community. We also make the harmonized data-sets with their metadata available to the wider research community.^2^

## 5 Conclusion

We have addressed the challenge of harmonizing large, and highly diverse public data for downstream meta-analysis. Aiming at high community acceptance and a computationally efficient solution, we extend the well-established ComBat algorithm through the addition of regularization. We evaluate our novel algorithm on simulated, and public microarray and bulkRNAsq data. A variety of evaluation metrics attest comparable, or superior correction of batch-effects as established baseline models. Our analysis constitutes a proof of principle to motivate and enable further large-scale meta-analyses.

## Footnotes

1 https://github.com/BorgwardtLab/*reComBat*

2 https://github.com/BorgwardtLab/batchCorrectionPublicData

