## Supplement for "reComBat: batch-effect removal in large-scale multi-source gene-expression data integration"

### A Evaluating batch-correction efficacy

We aim to provide an in-depth characterization of the harmonized data. An optimal input for meta-analysis comprises a range of metadata subsets with multiple batches for each of them. Rigorous, quantitative evaluation of different aspects of the correction based on metrics is key. Our analysis is based on collections of samples with the same experimental design. Inspired by the graph theoretical notion of n-hop neighborhoods [10], we group samples into so-called *Zero-Hops*. Each Zero-Hop defines a set of samples which share the exact same experimental design. Data harmonization efficacy can be quantified in terms of Zero-Hop purity and batch impurity. Note that the Zero-Hop label is used for evaluation purposes only and not as an input to *reComBat*.

We implemented seven evaluation metrics quantifying the sample distance measures, cluster (im-)purity and batch/design classification performance on the obtained transcription profiles. batch-correction methods were compared with respect to the following metrics with statistical significance being evaluated by Mann-Whitney U test.

##### Sample distance- and neighborhood-based metrics

*Cross-distances:* We calculate the median distance between all samples of a Zero-Hop and divide it by the median distance of all data points independent of batch or Zero-Hop. Smaller distances for Zero-Hops and large distances for batch are desired. The cross-distance does not account for distances of individual samples.

*Distance Ratio Score:* The distance ratio score (DRS) [14] quantifies the "closeness" of samples originating from the same condition versus the closeness of samples which should not. The DRS metric for a data-set of $n$ samples is defined as

$$\mathrm{DRS}_{\log} = \frac{1}{n}\sum_{i=1}^{n} \log_2\left(\frac{d(\boldsymbol{Y}_i, \boldsymbol{Y}_{i,dt})}{d(\boldsymbol{Y}_i, \boldsymbol{Y}_{i,db/st})}\right), \tag{1}$$

where $d(\cdot,\cdot)$ is a distance metric, $\boldsymbol{Y}_i$ is the $i^{\text{th}}$ sample and $\boldsymbol{Y}_{i,dt}$, $\boldsymbol{Y}_{i,db/st}$ are the closest samples from a different Zero-Hop (different type) and the same Zero-Hop (same type) but different batch respectively. A good correction of batch-effects results in an increase in $\mathrm{DRS}_{\log}$.

*Shannon Entropy:* Following [5], we calculated the local Shannon entropy with respect to batch and Zero-Hops within sets of $N$ nearest neighbours to a sample striving for high batch entropy while maintaining a small Zero-Hop entropy. For each sample $k$ the entropy $S$ with respect to batch $b$ is defined locally as

$$S_k = -\sum_{b=1}^{n_b} p_b \ln(p_b), \tag{2}$$

where $p_b$ denotes the locally estimated probabilities of the different batches and $n_b$ is the total number of batches. The Zero-Hop entropy is defined analogously. We chose the number of nearest neighbours for entropy calculation to be $N = 14$ corresponding to the median number of samples per Zero-Hop.

##### Cluster-based metrics

*Minimum Separation Number:* We define minimum separation number to be an integer quantifying the overlap of Zero-Hop clusters. Based on agglomerative clustering, initially, all samples occupy a single cluster. Then the number of clusters is increased in unit steps. The smallest number of clusters that assigns a Zero-Hop to at least two clusters, is defined as the minimum separation number for this Zero-Hop. We report mean values and standard deviations required to separate all Zero-Hops.

*Cluster purity:* We cluster the (corrected) expression data into $n_{\mathrm{ZH}}$ clusters, where $n_{\mathrm{ZH}}$ is the number of Zero-Hops. The purity of each cluster $c$, purity($c$), is calculated as the ratio of the number of samples of the dominant Zero-Hop in $c$, $n_{d,c}$, over the cluster size, $n_c$,

$$\text{purity}(c) = \frac{n_{d,c}}{n_c}. \tag{3}$$

A purity of 1 corresponds to a cluster comprising only samples from a single Zero-Hop. With decreasing values more conditions are represented in the cluster.

*Gini impurity:* The Gini impurity is a measure from decision tree learning [6, Section 9.2.3] and quantifies the probability of mislabelling a randomly chosen element of a cluster according to the label distribution of the cluster. Again, we create $n_{\mathrm{ZH}}$ clusters and use the Zero-Hop assignment as label. The output cluster impurities are given by assigning a label to each cluster chosen by a majority vote and calculating the fraction of majority labels in each cluster.

$$\text{gini}(c) = 1 - \sum_{i=1}^{n_{\mathrm{ZH}}} p_i^2 \quad \text{with } p_i = \frac{n_{i,c}}{n_c}, \tag{4}$$

where $n_{i,c}$ is the number of samples of Zero-Hop $i$ in cluster $c$.

#### Classifier-based metrics

*Linear Discriminant Analysis:* The Linear Discriminant Analysis (LDA) optimizes hyperplanes to maximally separate data points according to their labels (Zero-Hop) [6, Section 4.3]. We perform a stratified 10-fold cross validation and compare the LDA score (classification rate) on the held out test set. Mean values and standard deviations over the folds are reported.

*Logistic-regression-based evaluation:* This approach is inspired by the adversarial training of normAE [12]. Any classifier should not be able to predict the batch from the gene expression profile of well integrated data. Conversely, the prediction performance of the experimental design (Zero-Hop) should increase. To this end, we used two logistic regression (LR) classifiers, one for batch and one for Zero-Hops. Again, we perform a stratified 10-fold cross-validation and report the mean test set agglomerated balanced accuracy and F1-Scores.

### B Experimental data: data simulation and design annotation

#### B.1 Synthetic data

Since it is in general not possible to evaluate batch-correction methods on real-data due to the absence of a ground-truth we extensively tested our method on synthetic data. In particular, hyperparameters for *reComBat* which proved effective on synthetic data were carried over to the real-world data-sets. A further limitation to more testing of *reComBat* on real data is that such datasets are incredibly hard to generate as the design matrix has to be manually annotated by collating the relevant information by hand from the individual publications integrated into the dataset.

To simulate as realistic a dataset as possible, we assume that experimental data measured with identical experimental design and no batch effects follows a multivariate Gaussian distribution. We then shifted the data according to their experimental conditions by creating a design matrix this is singular. We introduced a number of design features and each feature can assume one of a number of categories, similar to the real data-sets we study. To induce singularity we simply make the last design feature dependent on the first, by evenly permuting its categories. Note that in principle dependent design features are an indicator of poor study design, however, when integrating real-world data-sets, such as our pseudomonas microarray data-set, this does indeed happen to be the case. Furthermore, additive and multiplicative batch effects were introduced. To this end we randomly shifted a separate set of 20% of the genes in each batch.

The classic train/test setup is not feasible in many real-world applications and common methods such as Harmony and scGen do not provide the option to test on a separate test set, even though in ComBat, respectively *reComBat*, this is possible. In general, train/test or cross-validation experiments are not common in the batch-correction literature due to their infeasibility.

Note that, as our method provides only a small, yet necessary, change to the original ComBat method [7], we would expect ComBat to perform similarly in the case of a non-singular design matrix.

We tested our method as a function of sample size, number of genes (to also highlight its efficacy in the case of high-dimensional RNAsq experiments), number of samples-per-batch, number of zero-hops, and relative batch-effect strength. The methods were evaluated using the LDA score, cluster impurity, and entropy scores. The runtime of the methods was also studied, noting that in any scenario, the (parametric) reComBat methods was extremely fast.

#### B.2 Microarray data

Data was collected from the GEO database [3] (accessed October 2020). All entries on PA using the GPL84 Affymetrix GeneChip were considered. The GPL84 array comprises a total of 5900 probe sets including annotated genome of PA01 (5,568 genes) and other PA strains (117 genes), as well as 199 intergenic regions. In total, $n = 1260$ samples within 150 independent batches (GSE identifiers) were identified. The expression data were subjected to Robust Multi-array Averaging (RMA) using the *justRMA* function from the *affy* *R*-package. The relevant experimental design regarding culture conditions and PA strain was extracted manually and coarsened as outlined in Appendix A. For quality control (QC) we deleted all samples with incomplete design information and single-sample batches. Due to the large quantity of unique modifications, we dropped additional information regarding genetic alterations (mutations, plasmids, etc.) as part of the QC. As a final step of the preprocessing, we pruned all single-batch Zero-Hops. A detailed overview of all design categories contributing to the Zero-Hop definition is given in Appendix A. Given the the imposed exclusion criteria, we analysed a total of 887 samples, structured within 39 Zero-Hops from 114 individual GSEs (i.e. batches) comprising 5 to 170 individual experiments each (see Table S1).

#### B.3 bulkRNAsq data

RNAsq read data was collected from the GEO, SRA and ENA data bases (accessed February 2021). All published data sets using the PAO1 and PA14 reference strains, with more than two replicates were considered. Data was preprocessed using the Galaxy Project [1]. Briefly, read quality was ensured (minimum base quality of 30, minimum length of 40 bases per read) and, when appropriate, sequencing adaptors were removed using trimmomatic v0.38 [4]. Reads were aligned using HISAT2 version 2.2.1 [8], using the PAO1 or

PA14 genomes when appropriate (accession numbers AE004091.2 and NC_008463.1). The number of reads mapping to each feature was calculated using htseq-count version 0.9.1 [2]. 5292 unique PAO1 and PA14 orthologs were matched using OrthologeDB [15]. In total, $n = 340$ samples within 32 independent batches (GSEs) were identified with coarsened meta-information and assigned to 12 Zero-Hops (see Table S2)). The obtained design matrix is not-singular, however, ill-conditioned.

##### B.4 Zero-Hop assignment

Each sample (independent of bulkRNA or micro array data) was assigned to one of the available categories for each of the following design categories:

- **Temperature** (three conditions): room temperature (20-25°C), body surface temperature (28-32°C), body temperature (36-38°C). In case of missing data, body temperature was assumed.
- **Growth phase** (three conditions): lag phase (optical density below or equal to 0.2), exponential growth, plateau (growth time > 7h, or optical density $\geq$2.5) - exponential growth was assumed in case of missing information for *in vitro* cultures.
- **Culture medium** (five conditions): poor (no specific growth promoting substrate, e.g. water), defined (generally minimal media, e.g. M9 Minimal Salts), rich (nutrient rich media, e.g. Lysogeny broth (LB)), in vivo-like (e.g. artificial urine) and *in vivo* (both human and animal models). For subsets of defined media we further added additional description regarding the medias' gluconeogenic properties (yes, no, both, undefined). For 'rich' media such as LB, we further accounted for a relative richness score summarising the nutritional complexity of the media. Said score was determined using LB as rich medium reference. Any media that contained less nutrients than LB (such as diluted LB or minimal media) was classed as "less rich" and any media that contained higher nutrient concentrations was classed as "richer" (such as terrific broth (TB)).
- **Culture substrate** (four conditions): liquid, film, plate, *in vivo*. Imputation of missing data with 'liquid'.
- **Oxygenation** (two conditions): hypoxic, aerobic. Aerobic growth conditions were assumed to be the standard.
- **PA Strain** (twenty conditions): We included data on the laboratory strains PAO1, PA1161, H103, PA14, MPAO1,DK2, PAO25, ATCC 33988, PAO1161, PAO1ut, PAO6261, UCBPP-PA14, TBCF10839, PAK, PA2206, UUPA85, UUPA38, NCTC8626, PAHM4, as well as "Clinical Isolates" or "Uncharacterized" for unspecified clinical samples.
- **Antibiotic exposure** (three conditions): no exposure (no ABX), interval treatment (treat), continuous exposure (consistent).

An overview of the specific experimental designs for all Zero-Hops is shown in Tables S1 and S2.

| **Zero-Hop** | **Strain** | **Growth phase** | **Culture** | **Temperature** | **Oxygenation** | **Medium** | **Antibiotic** | $n_{Samples}$ | $n_{Batches}$ |
|---|---|---|---|---|---|---|---|---|---|
| **0** | PAO1 | Exp | liquid | body temp. | hypoxic | rich | no ABX | 48 | 2 |
| **1** | PAO1 | Plat | film | body temp. | aerobic | less rich | no ABX | 12 | 2 |
| **2** | PAO1 | Exp | liquid | body temp. | hypoxic | defined | no ABX | 10 | 2 |
| **3** | PAO25 | Exp | liquid | body temp. | aerobicc | rich | consistent | 10 | 3 |
| **4** | PAO14 | Exp | liquid | body temp. | aerobic | rich | treat | 11 | 2 |
| **5** | PAO1 | Plat | film | room temp. | aerobic | defined | no ABX | 9 | 3 |
| **6** | PAO1 | Plat | liquid | body temp. | aerobic | rich | treat | 9 | 4 |
| **7** | PAO14 | Plat | film | body temp. | aerobic | rich | no ABX | 15 | 5 |
| **8** | PAO1 | Exp | liquid | surface temp. | aerobic | rich | no ABX | 7 | 2 |
| **9** | PAO1 | Plat | liquid | body temp. | hypoxic | rich | no ABX | 11 | 2 |
| **10** | PAO1 | Plat | liquid | body temp. | aerobic | less rich | no ABX | 9 | 2 |
| **11** | PAO1 | Exp | liquid | body temp. | aerobic | less rich | no ABX | 21 | 4 |
| **12** | PAO1ut | Exp | liquid | body temp. | aerobic | rich | no ABX | 8 | 2 |
| **13** | PAO1 | Exp | film | body temp. | aerobic | rich | no ABX | 7 | 2 |
| **14** | PAO1 | Plat | plate | body temp. | aerobic | rich | no ABX | 18 | 3 |
| **15** | PAO1 | Plat | film | body temp. | aerobic | rich(0) | treat | 8 | 3 |
| **16** | Clinical isolate | Exp | liquid | body temp. | aerobic | in vivo like | no ABX | 114 | 7 |
| **17** | PAO1 | Exp | liquid | body temp. | aerobic | rich | no ABX | 5 | 2 |
| **18** | TBCCF10839 | Exp | liquid | body temp. | aerobic | defined | no ABX | 14 | 2 |
| **19** | PAO14 | Exp | liquid | body temp. | aerobic | rich | no ABX | 12 | 2 |
| **20** | PAO1 | Exp | liquid | body temp. | aerobic | rich(0) | treat | 24 | 5 |
| **21** | PAO1 | Plat | film | body temp. | aerobic | in vivo like | no ABX | 6 | 2 |
| **22** | PAO14 | Plat | liquid | body temp. | aerobic | rich | no ABX | 20 | 4 |
| **23** | PAO1 | Plat | film | body temp. | aerobic | defined | treat | 10 | 2 |
| **24** | Clinical isolate | in vivo | liquid | body temp. | aerobic | in vivo | no ABX | 12 | 2 |
| **25** | Clinical isolate | Plat | film | body temp. | aerobic | rich | no ABX | 16 | 3 |
| **26** | PAO1 | Exp | liquid | body temp. | aerobic | gluconeogenic | no ABX | 48 | 6 |
| **27** | PAO14 | Exp | liquid | body temp. | aerobic | rich | no ABX | 14 | 5 |
| **28** | MPAO1 | Exp | liquid | body temp. | aerobic | defined | no ABX | 7 | 2 |
| **29** | PAO1 | Plat | liquid | body temp. | aerobic | defined | no ABX | 24 | 3 |
| **30** | PAO1 | Plat | film | body temp. | aerobic | defined | no ABX | 24 | 4 |
| **31** | PAO1 | Plat | liquid | surface temp. | aerobic | rich | no ABX | 14 | 3 |
| **32** | PAO1 | Plat | film | body temp. | aerobic | rich(0) | no ABX | 22 | 6 |
| **33** | PAO1 | Plat | liquid | body temp. | aerobic | rich(0) | no ABX | 38 | 9 |
| **34** | PAO1 | Exp | liquid | body temp. | aerobic | richer | no ABX | 20 | 3 |
| **35** | PAO1 | Exp | liquid | body temp. | aerobic | rich(0) | no ABX | 170 | 23 |
| **36** | PAO1 | Exp | liquid | body temp. | aerobic | (not)gluconeogenic | no ABX | 10 | 2 |
| **37** | PAO1 | Exp | liquid | body temp. | aerobic | not defined | no ABX | 24 | 5 |
| **38** | PAO1 | Exp | liquid | body temp. | aerobic | not gluconeogenic | no ABX | 26 | 4 |

**Table S1.** Overview of the included micro array data obtained from the GEO database with manually assigned coarse design groups for each Zero-Hop. The number of samples $n_{Samples}$ and batches $n_{Batches}$in each Zero-Hop are also given. Note, that batches may overlap between the Zero-Hops. Abbreviations: Plat: Plateau phase, Exp: exponential growth phase, temp: temperature, ABX: antibiotic treatment, consistent: continuous antibiotic treatment, treat: interval antibiotic treatment

| **Zero-Hop** | **Strain** | **Growth phase** | **Culture** | **Temperature** | **Oxygenation** | **Medium** | **Antibiotic** | $n_{Samples}$ | $n_{Batches}$ |
|---|---|---|---|---|---|---|---|---|---|
| **0** | Clinical isolate | Plat | liquid | body temp. | aerobic | rich | no ABX | 7 | 2 |
| **1** | PAO1 | unknown | liquid | body temp. | aerobic | less rich | no ABX | 12 | 2 |
| **2** | PAO14 | Exp | liquid | body temp. | aerobic | rich | no ABX | 27 | 4 |
| **3** | PAO1 | unknown | in vivo | body temp. | aerobicc | in vivo | no ABX | 7 | 2 |
| **4** | PAO14 | Exp | liquid | body temp. | aerobic | rich | no ABX | 42 | 3 |
| **5** | PAO1 | unknown | liquid | body temp. | aerobic | in vivo like | no ABX | 3 | 2 |
| **6** | PAO1 | unknown | liquid | body temp. | aerobic | defined | no ABX | 10 | 2 |
| **7** | PAO14 | unknown | in vivo | body temp. | aerobic | in vivo | no ABX | 35 | 3 |
| **8** | PAO14 | Exp | liquid | body temp. | aerobic | defined | no ABX | 44 | 4 |
| **9** | PAO14 | Plat | liquid | body temp. | aerobic | rich | no ABX | 17 | 2 |
| **10** | PAO1 | Exp | liquid | body temp. | aerobic | rich | no ABX | 126 | 15 |
| **11** | PAO1 | Plat | liquid | body temp. | aerobic | rich | no ABX | 10 | 2 |

**Table S2.** Overview of the included bulkRNAsq data obtained from the GEO, SRA and ENA databases with the manually assigned coarse design groups for each Zero-Hop. The relevant number of samples $n_{Samples}$ and batches $n_{Batches}$ are also given. Abbreviations: Plat: Plateau phase, Exp: exponential growth phase, temp: temperature, ABX: antibiotic treatment.

#### C Baseline methods

As baselines we chose representative examples from location-scale, matrix factorization, feature-based and deep learning. Unless mentioned otherwise, all methods employ the full design matrix as covariates, Zero-Hop information was only used for assessment rather than model-fitting.

*Standardization:* For each sample we calculated the mean and standard deviation over all genes and Z-scored the expression profiles accordingly. All subsequent methods took the standardized data as input.

*Marker Gene elimination:* We eliminated the top $n$ marker genes of each batch pair from the data. The optimal number of genes to be eliminated (eight) was identified by grid search. This reduced the features to 5150/5048 out of 5900/5292 genes for microarray/bulkRNAsq data.

*Principal component elimination:* The first $n$ principal components (PCs) of each Zero-Hop were calculated, and PCs accounting for more than $k\%$ of the variance explained were subtracted from the expression matrix. We identified $n = 20\%$ of the number of samples in the Zero-Hop, but minimum three, and a cut-off explained variance of 10% to be optimal by grid search.

*Combat:* As discussed above the solution of the ordinary linear regression problem in case of a singular design matrix is not unique and hence standard python implementations of ComBat based on matrix inversion fail. We implemented a version which does not solely rely on matrix inversion to handle such singular problems.

*Harmony:* Harmony [9] was previously suggested as the best practice method for single cell RNAsq batch-correction in a large-scale review [13]. It accounts for desired and unwanted variation by projecting samples to a shared low-dimensional embedding followed by soft k-means clustering penalizing low batch variety within clusters. Iteratively, cluster specific linear correction factors are calculated and each sample is eventually corrected by the weighted average over all corrections factors for the clusters it is a part of. We used the python *harmony-pytorch* package with default parameters.

*scGen:* scGen [11] combines variational autoencoders and latent space vector arithmetics. It previously out-performed other nonlinear batch-correction methods based on style-transfer Generative Adversarial Networks or conditional variational autoencoders. We used the python package *scgen* with default hyperparameters trained within 80-130 epochs with early stopping using all categorical covariates on a single NVIDIA TITAN RTX, 24 GiB RAM GPU. We also present results for scGen using the Zero-Hop as optimization label (denoted as scGen(Zero-Hop)). As such, scGen(Zero-Hop) specifically uses the Zero-Hop label (used for evaluation) as an input!

#### D reComBat hyperparameter selection

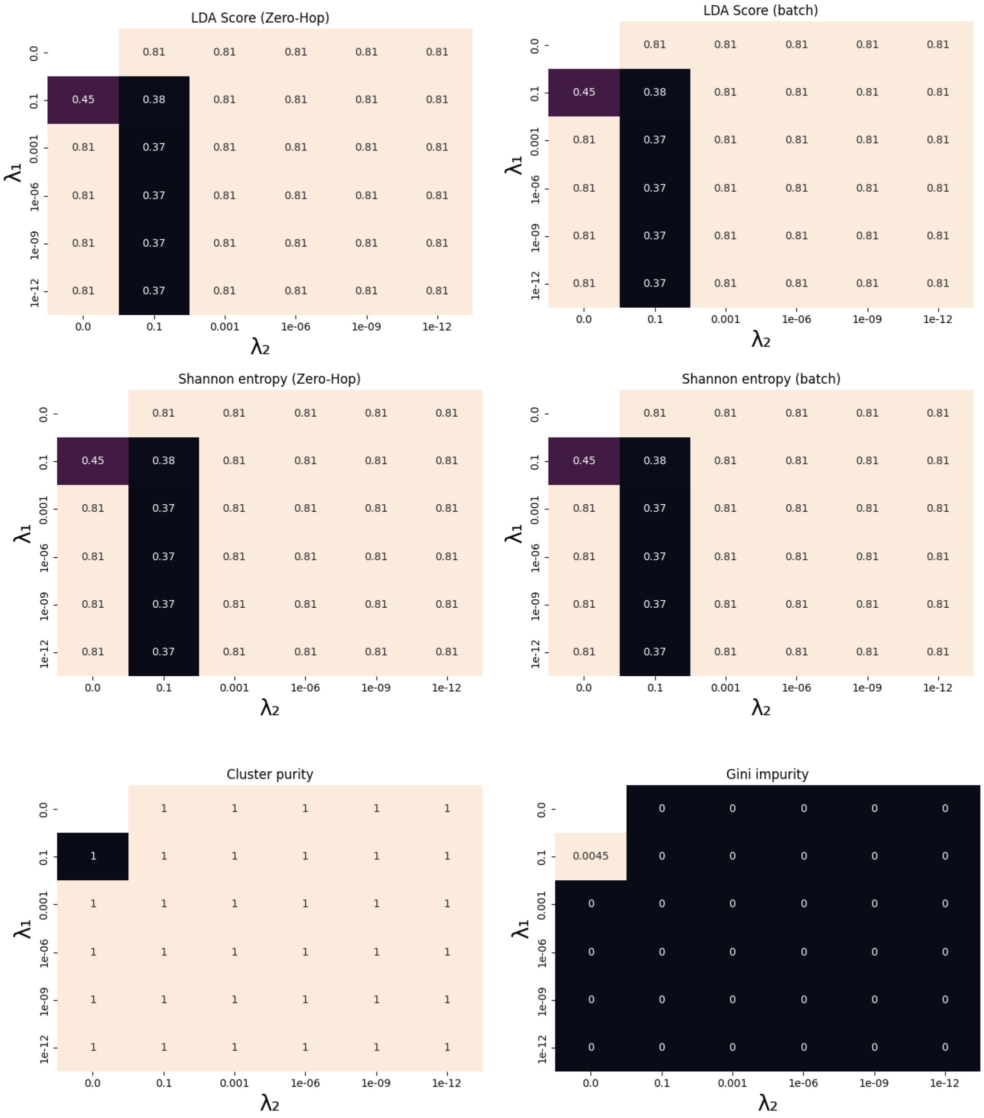


**Fig. S1.** Overview over LDA score, Shannon entropy, Gini impurity and cluster purity shown as heat-maps as a function of the regularization strengths $\lambda_1$ and $\lambda_2$ evaluated on synthetic data (6000 genes, 1000 samples, 30 Zero-Hops, 10 samples per batch). Mean values over 10 repeat runs are shown. Note that reComBat without regularization yielded numerically unstable results (i.e. nans).

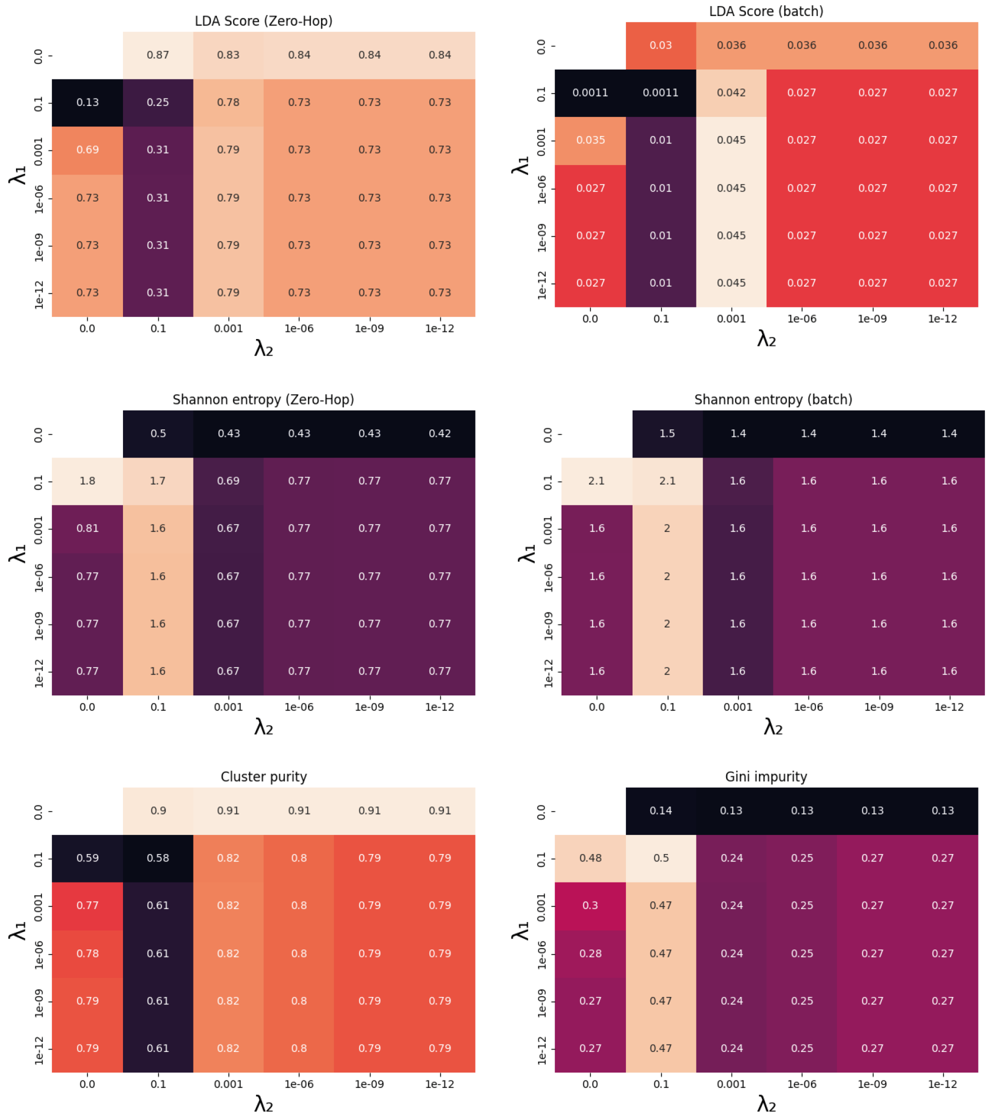


**Fig. S2.** Overview over LDA score, Shannon entropy, Gini impurity and cluster purity shown as heat-maps as a function of the regularization strengths $\lambda_1$ and $\lambda_2$ evaluated on microarray data. Mean values over 10 repeat runs are shown. Note that reComBat without regularization yielded numerically unstable results (i.e. nans).

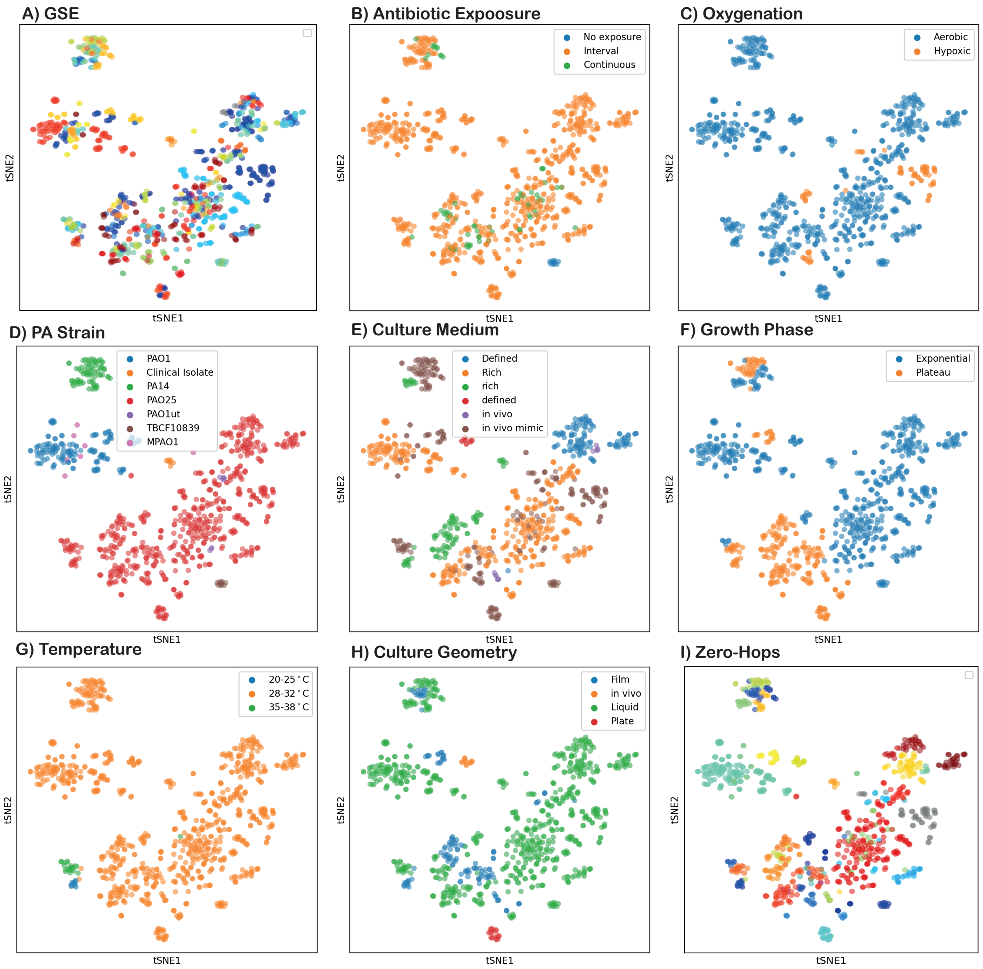


**Fig. S3.** tSNE plots of the corrected microarray data using ridge *reComBat* with a regularization strength $\lambda_2 = 1$ colored by batches (i.e. GSE) (A), all evaluated experimental designs (B-H), and the defined Zero-Hops (I).

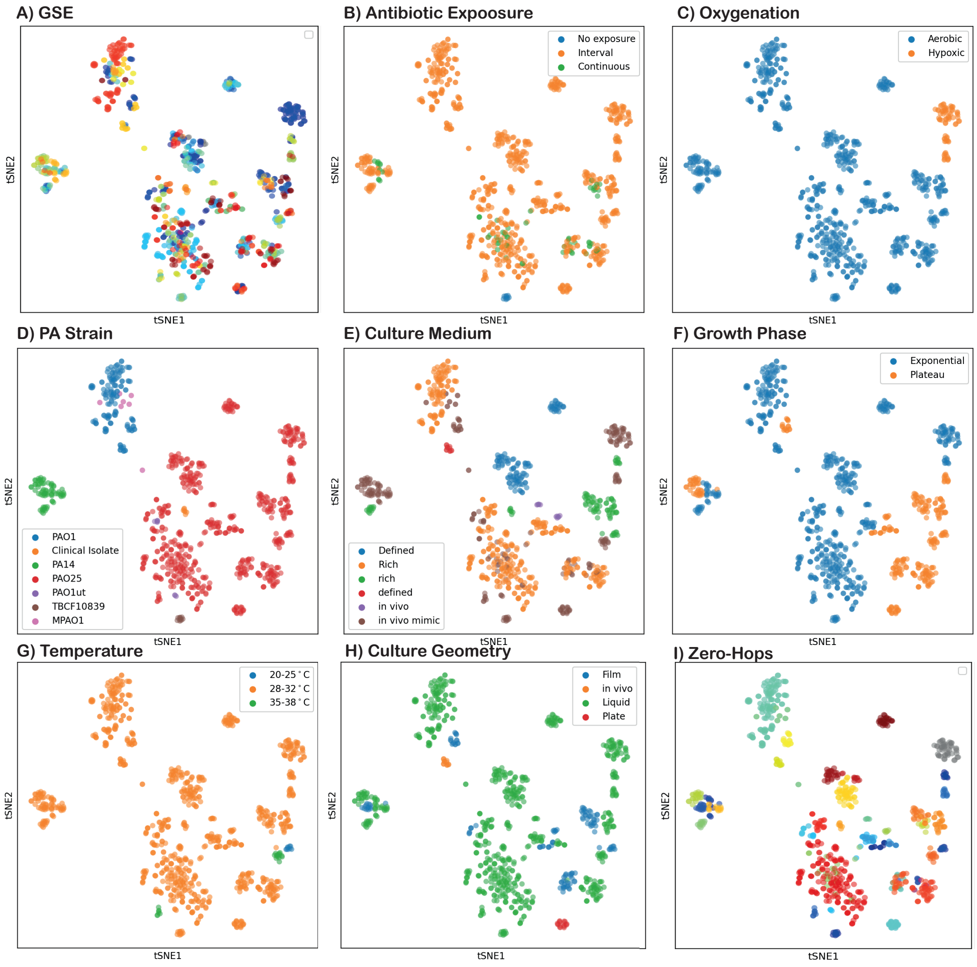


**Fig. S4.** tSNE plots of the corrected microarray data using ridge *reComBat* with a regularization strength $\lambda_2 = 0.1$ colored by batches (i.e GSE) (A), all evaluated experimental designs (B-H), and the defined Zero-Hops (I).

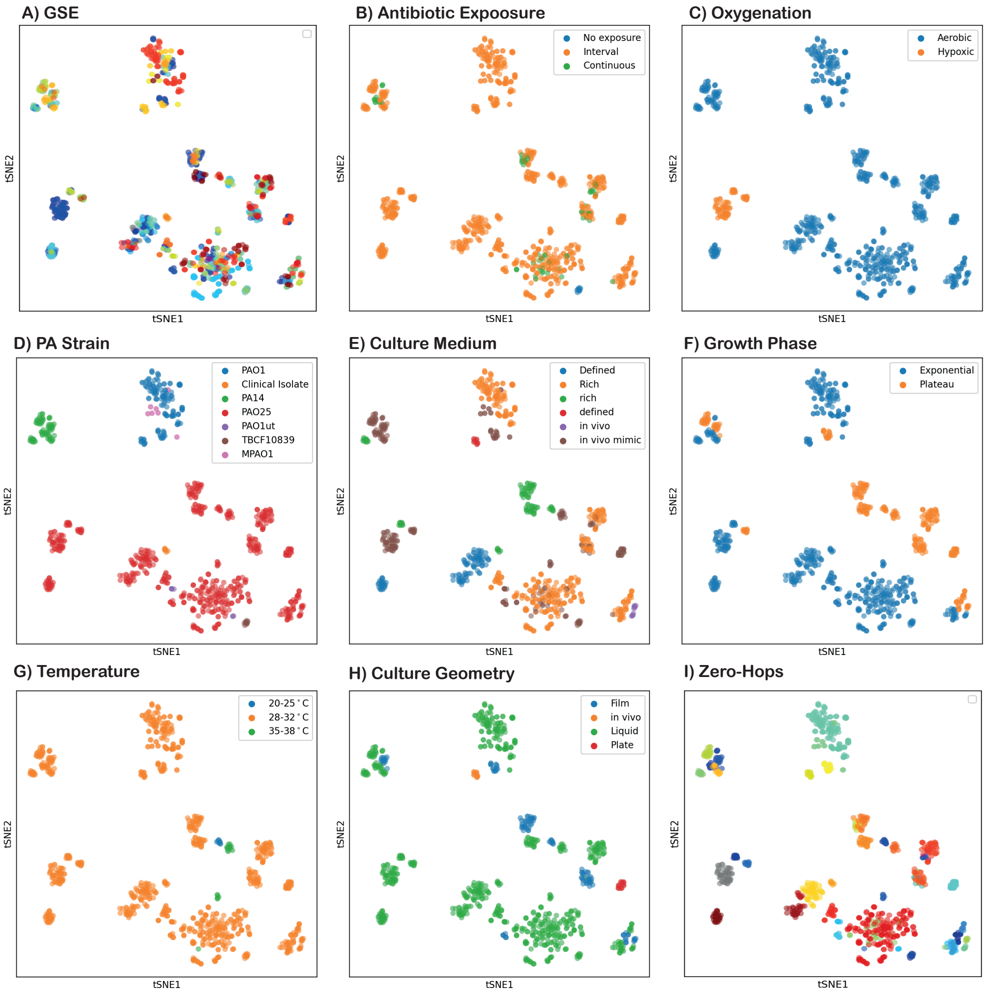


**Fig. S5.** tSNE plots of the corrected microarray data using ridge *reComBat* with a regularization strength $\lambda_2 = 1e{-}9$ colored by batches (i.e. GSE) (A), all evaluated experimental designs (B-H), and the defined Zero-Hops (I).

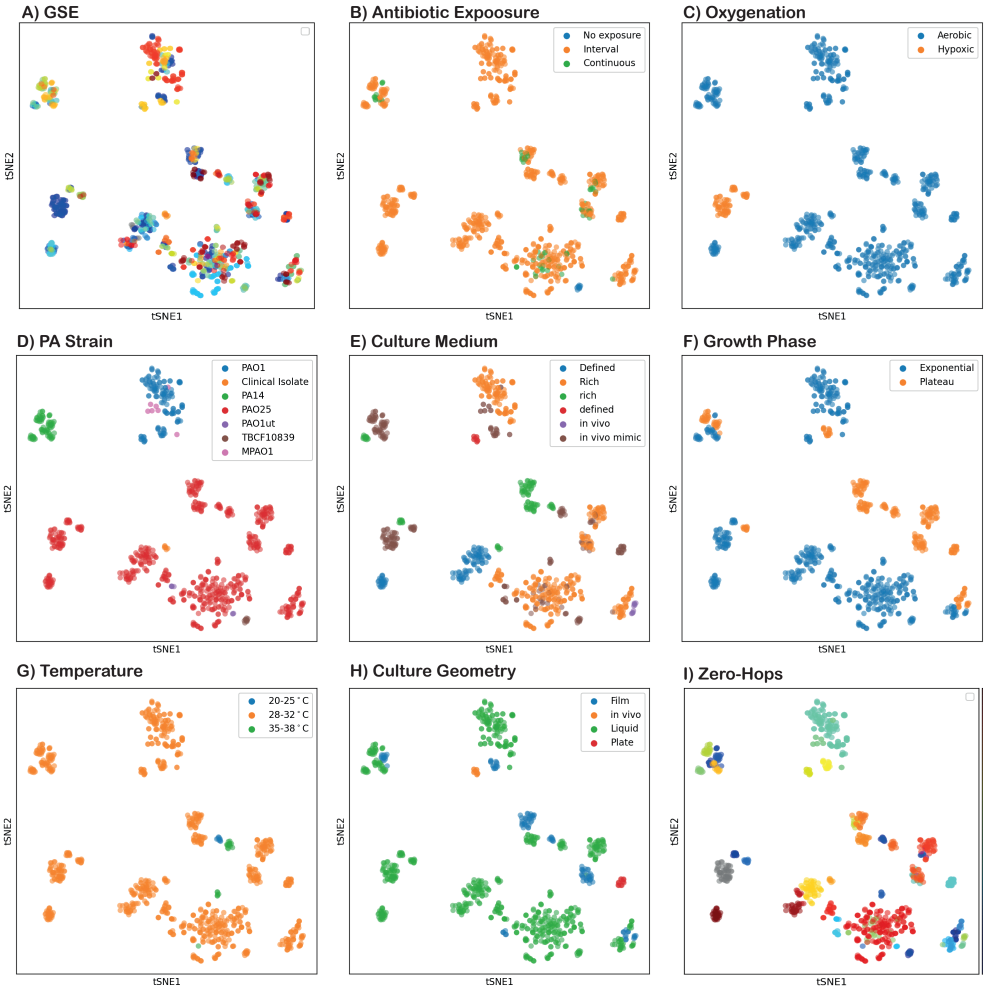


**Fig. S6.** tSNE plots of the corrected microarray data using LASSO *reComBat* with a regularization strength $\lambda_1 = 0.01$ colored by batches (A), all evaluated experimental designs (B-H), and the defined Zero-Hops (I).

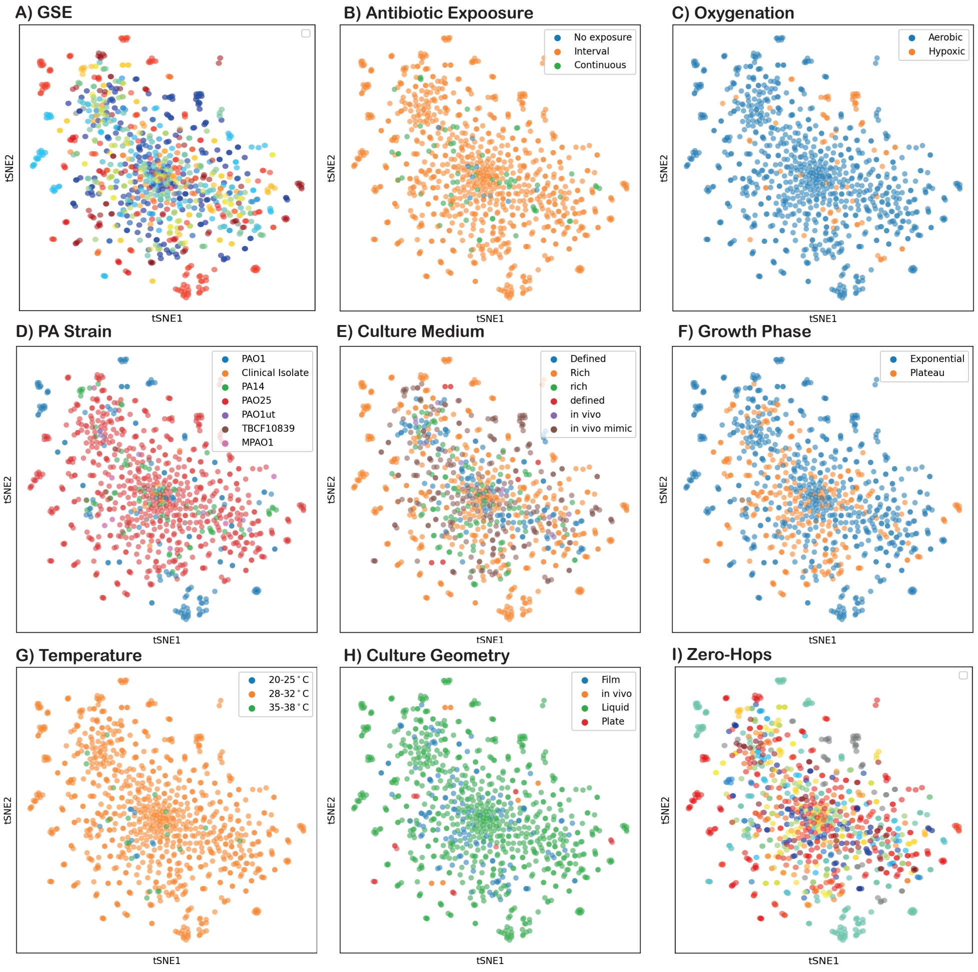


**Fig. S7.** tSNE plots of the corrected microarray data using *reComBat* with a regularization strength $\lambda_1 = 0.01$ and $\lambda_2 = 1e-9$ colored by batches (A), all evaluated experimental designs (B-H), and the defined Zero-Hops (I).

183 # E Additional results on synthetic data

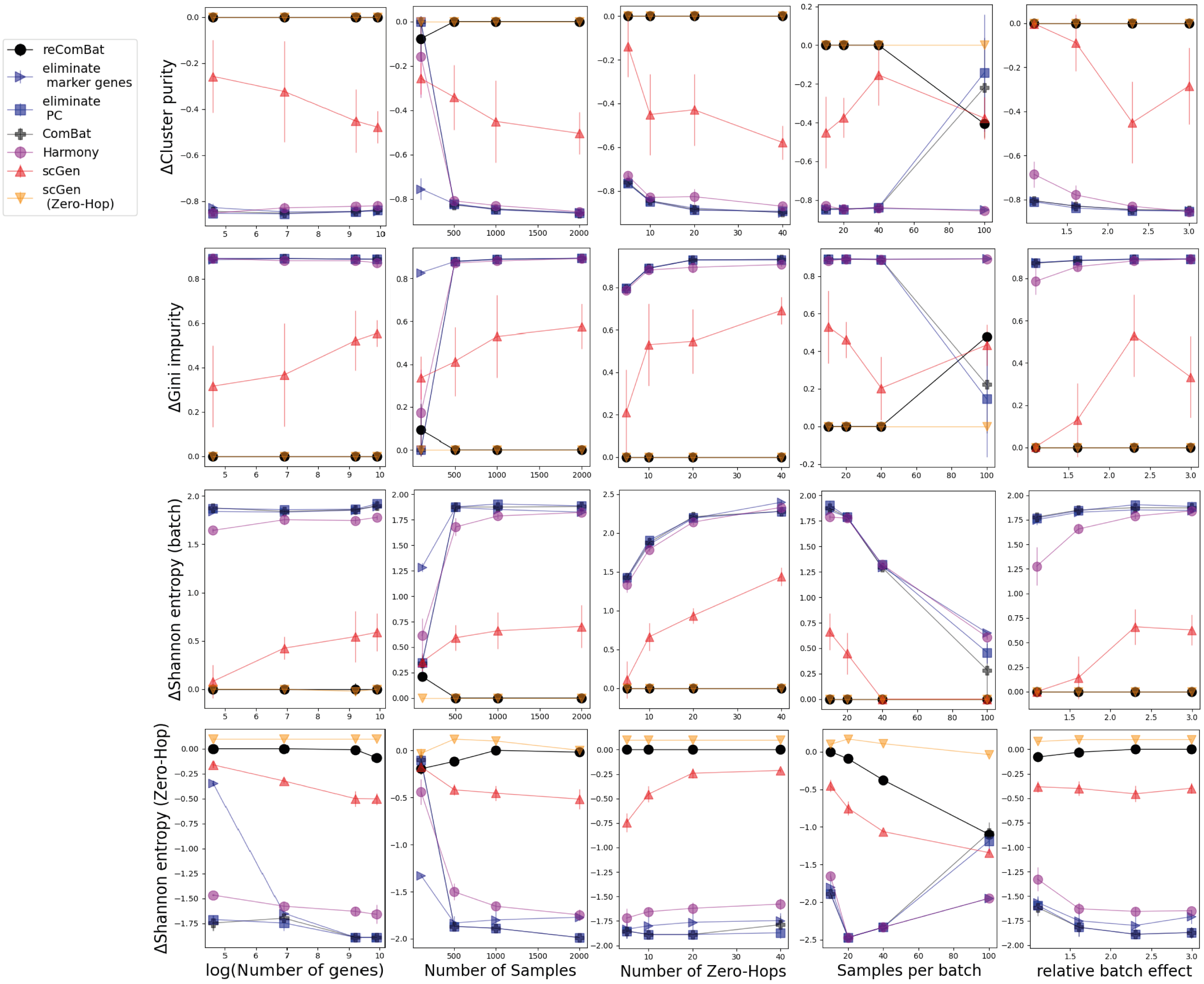


**Fig. S8.** Evaluation of simulated data as a function of the number of genes, samples, Zero-Hops, samples per batch and the disturbance strength of batch relative to metadata. Results are given as mean values and standard deviations for cluster purity (top), Gini impurity (2nd row), and Shannon entropy by batch (3rd row) or Zero-Hop(bottom).

### F tSNE plots for all (corrected) real-world data

#### F.1 Microarray data

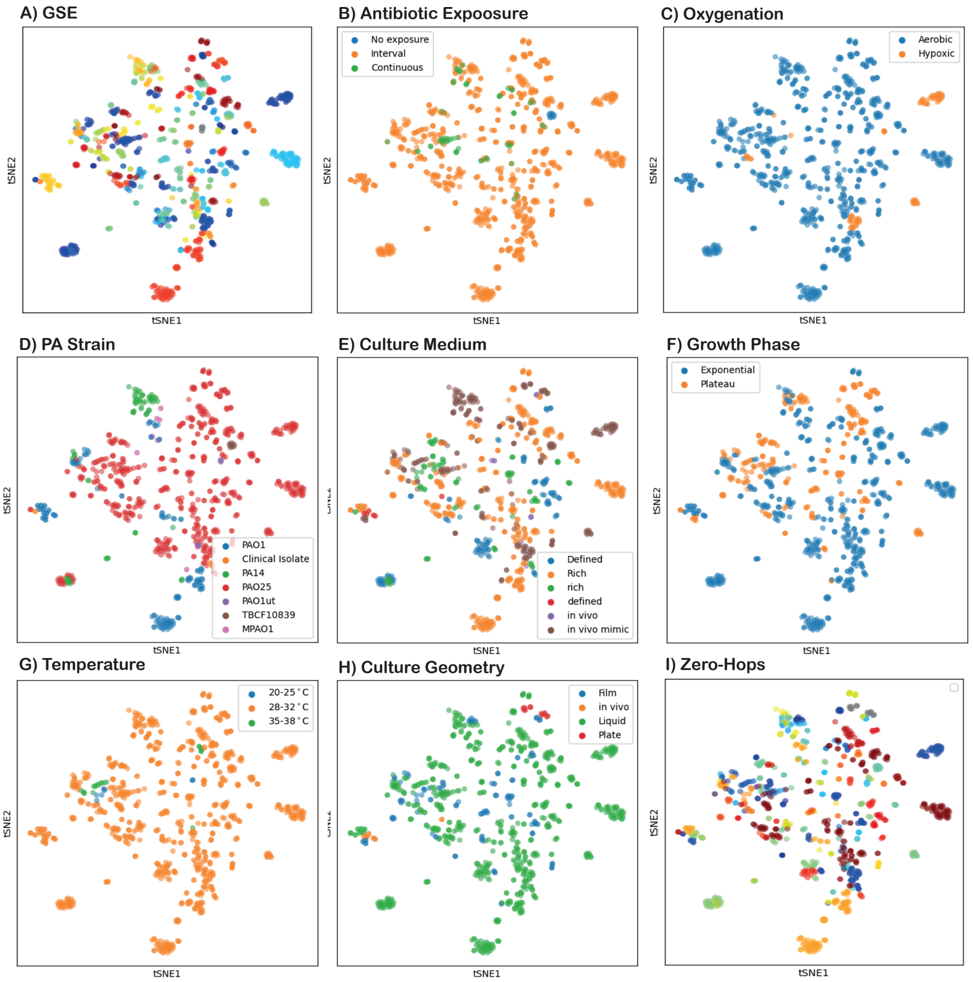


**Fig. S9.** tSNE plots of the uncorrected microarray data coloured by batches (A), all evaluated experimental designs (B-H), and the defined Zero-Hops (I). Clustering is largely driven by the GSE (i.e. batch) rather than by the underlying culture conditions or microbial strains.

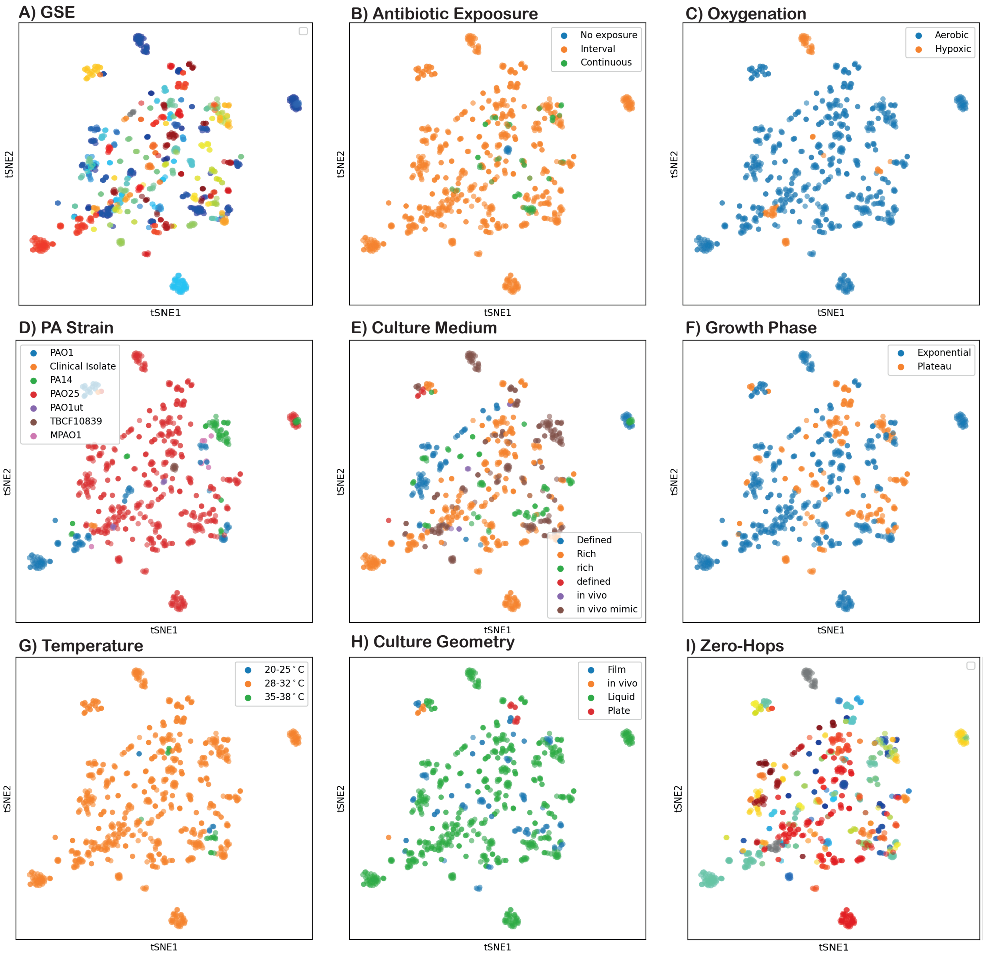


**Fig. S10.** tSNE plots of the corrected microarray data using Z-scoring (i.e. standardization) colored by batches (i.e. GSE) (A), all evaluated experimental designs (B-H), and the defined Zero-Hops (I).

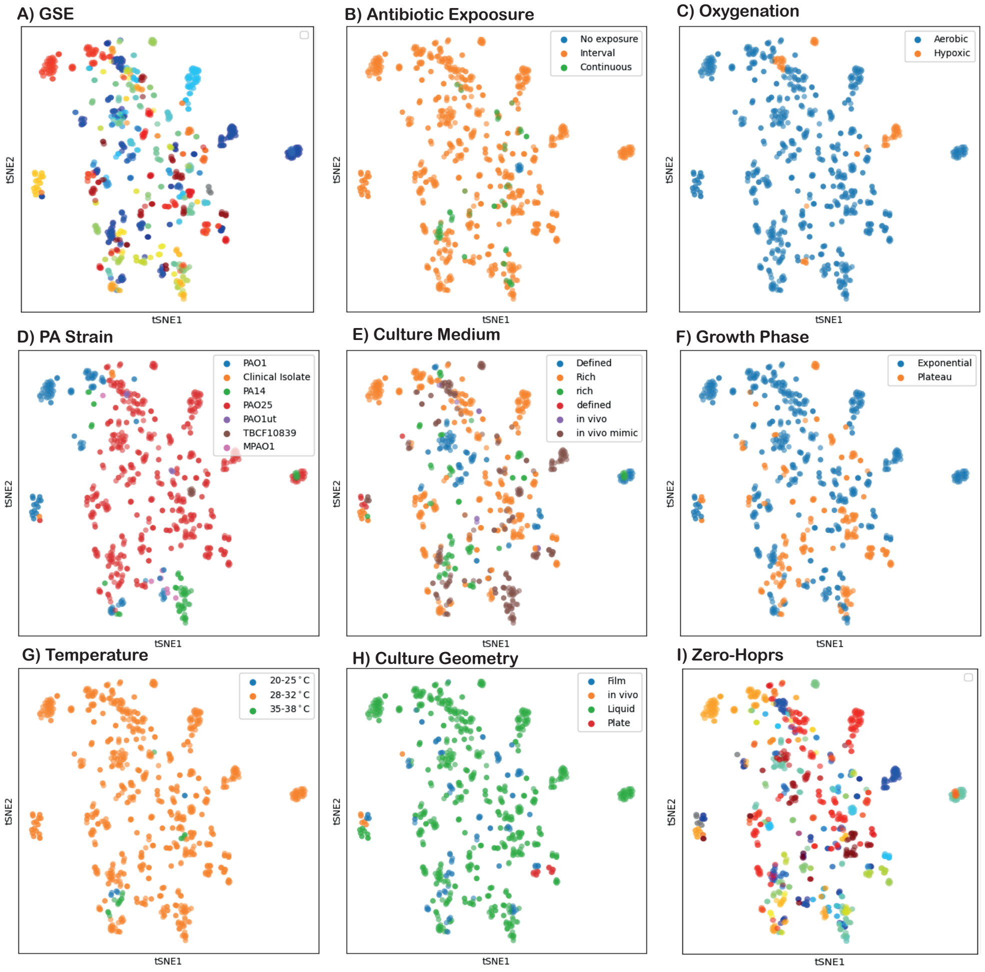


**Fig. S11.** tSNE plots of the corrected microarray data using marker gene elimination colored by batches (i.e. GSE)(A), all evaluated experimental designs (B-H), and the defined Zero-Hops (I).

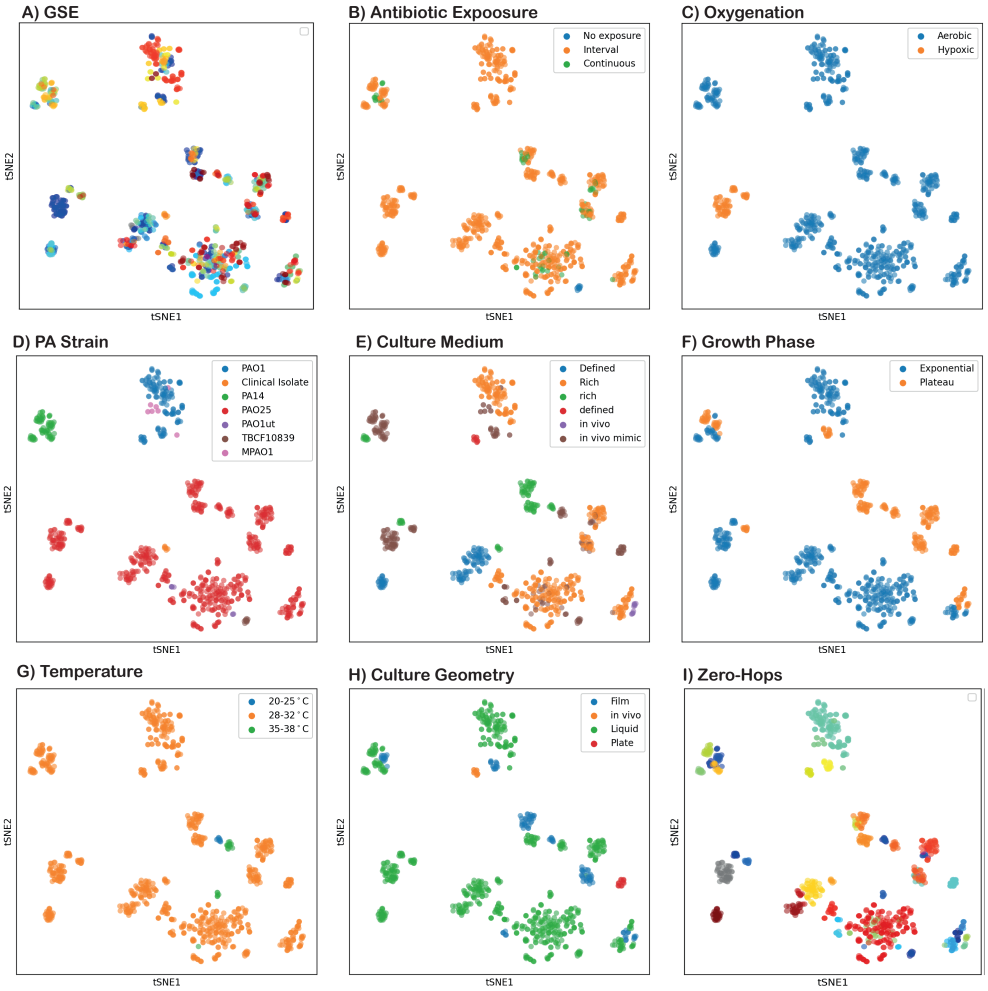


**Fig. S12.** tSNE plots of the corrected microarray data using PC elimination colored by batches (i.e. GSE)(A), all evaluated experimental designs (B-H), and the defined Zero-Hops (I).

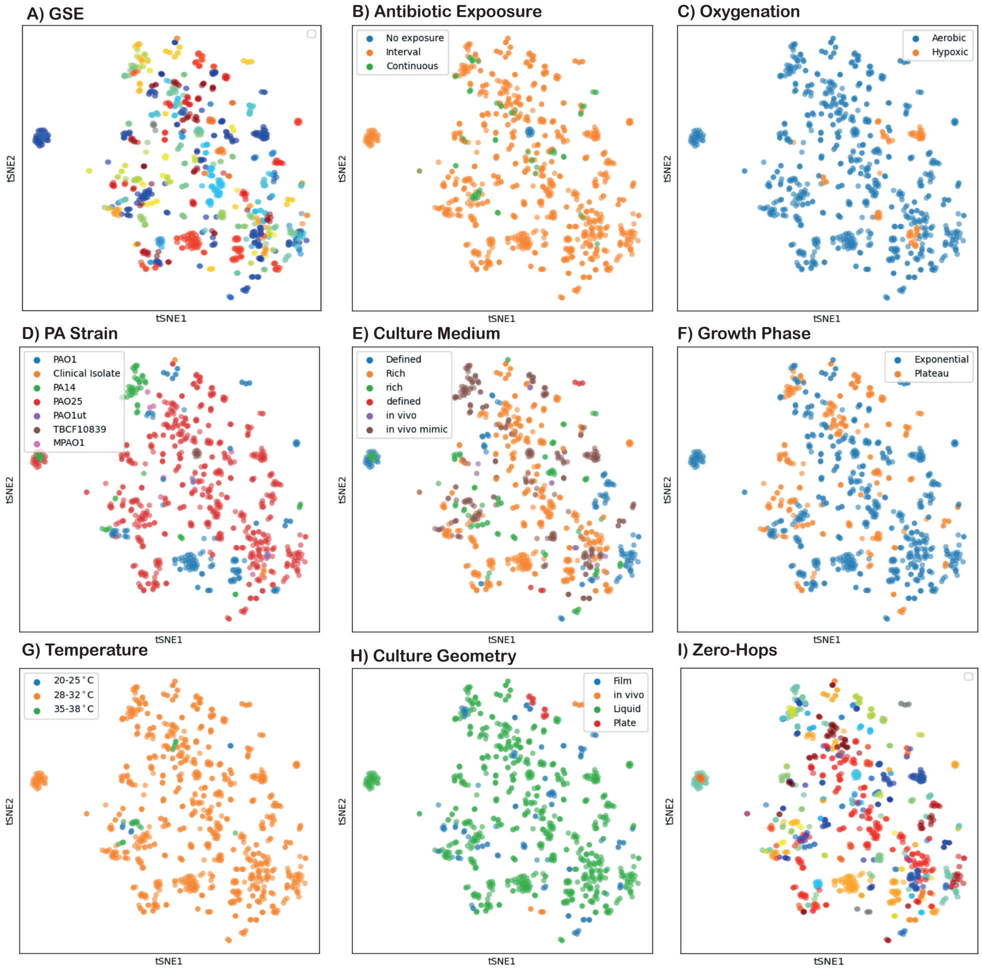


**Fig. S13.** tSNE plots of the corrected microarray data using Harmony colored by batches (i.e. GSE)(A), all evaluated experimental designs (B-H), and the defined Zero-Hops (I).

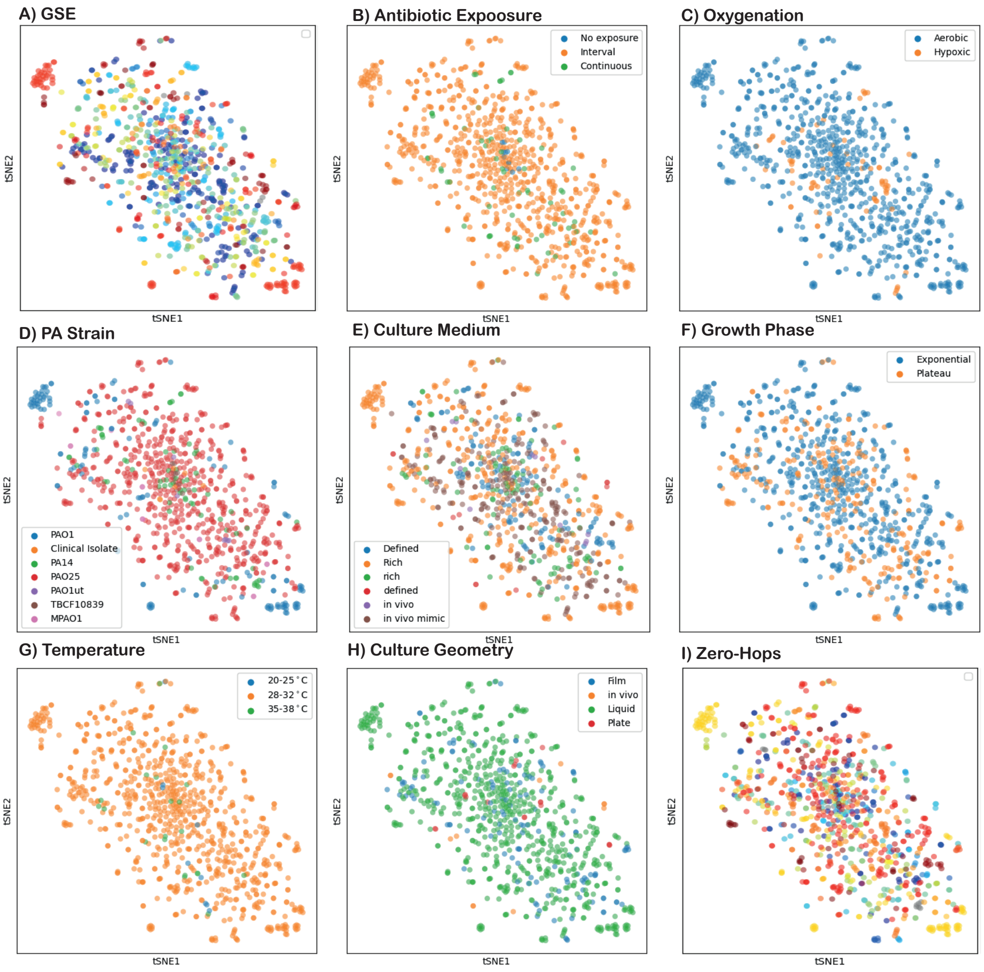


**Fig. S14.** tSNE plots of the corrected microarray data using scGen without information on Zero-Hops. Plots are colored by batches (i.e. GSE)(A), all evaluated experimental designs (B-H), and the defined Zero-Hops (I).

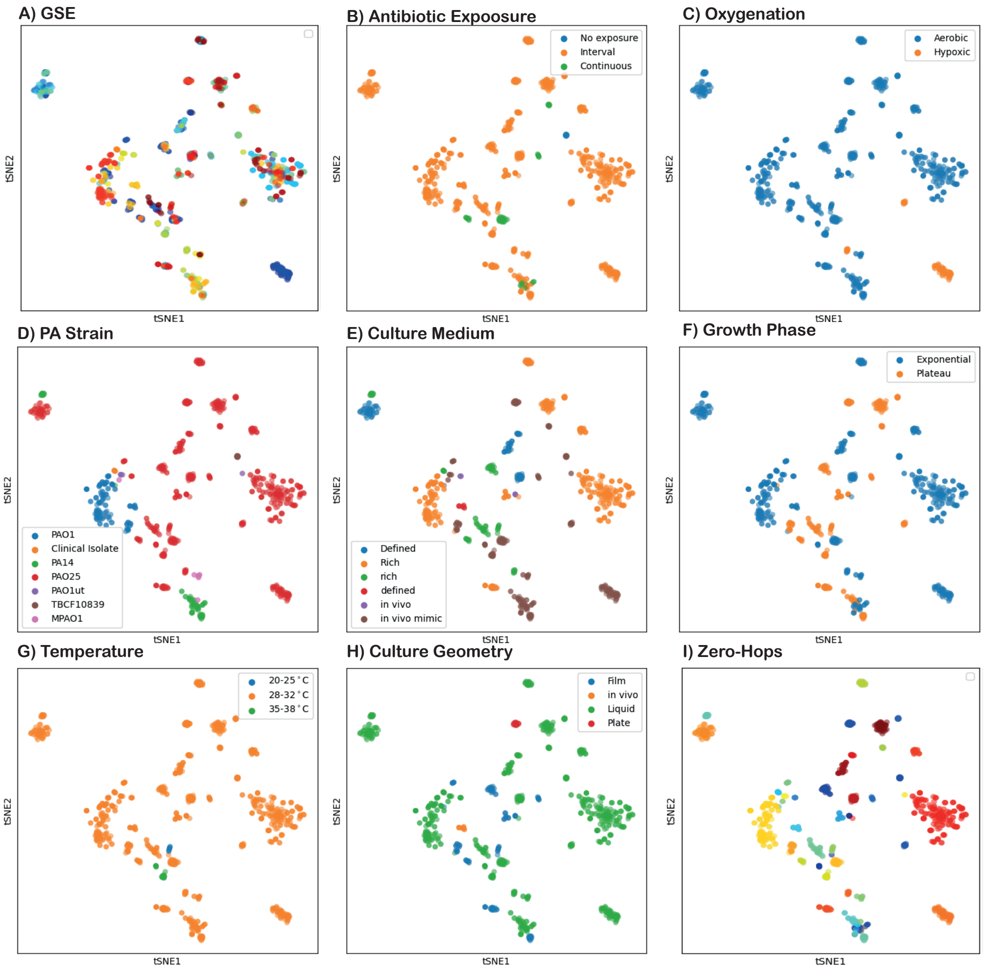


**Fig. S15.** tSNE plots of the corrected microarray data using scGen with Zero-Hop used as label. Plots are colored by batches (i.e. GSE)(A), all evaluated experimental designs (B-H), and the defined Zero-Hops (I).

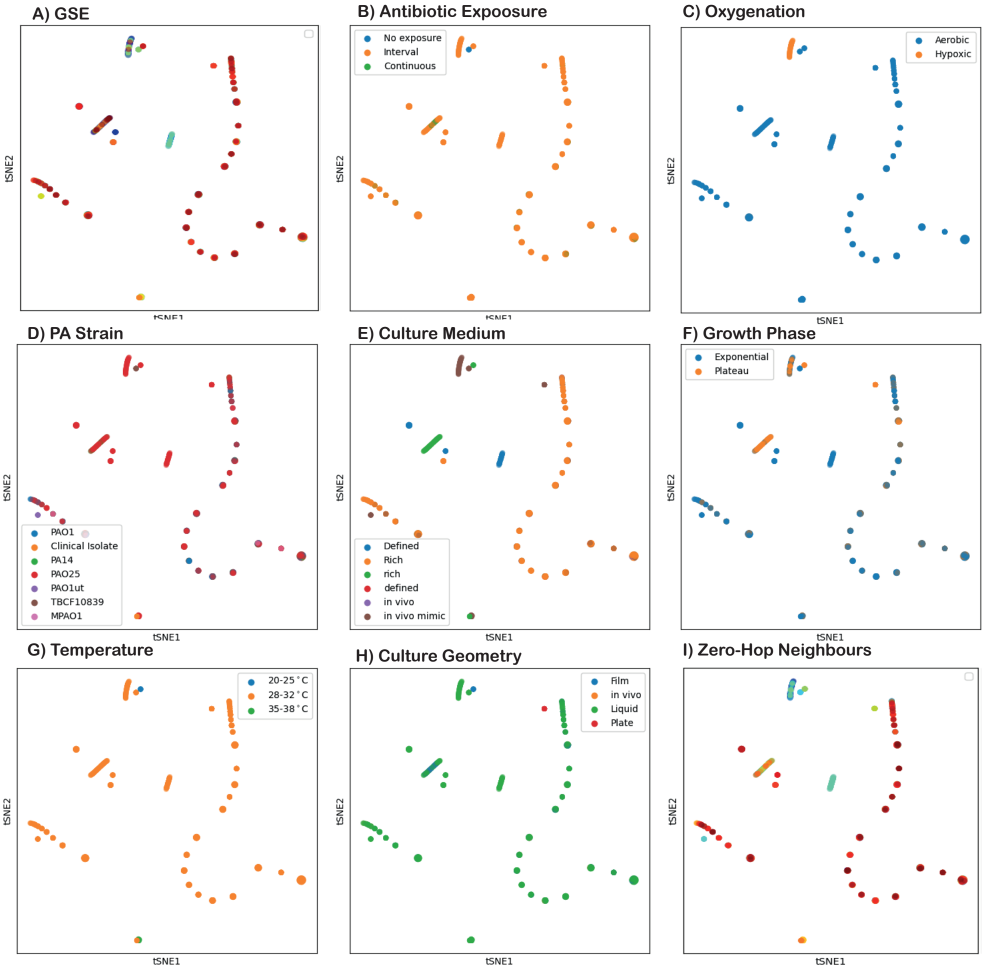


**Fig. S16.** tSNE plots of the corrected micro array data using ComBat. This correction did not yield a unique solution. Plots are colored by batches (i.e. GSE)(A), all evaluated experimental designs (B-H), and the defined Zero-Hops (I).

#### F.2 bulkRNAsq data

No batches evaluating different strains were available, leading to fairly clear separation of strains in uncorrected data.

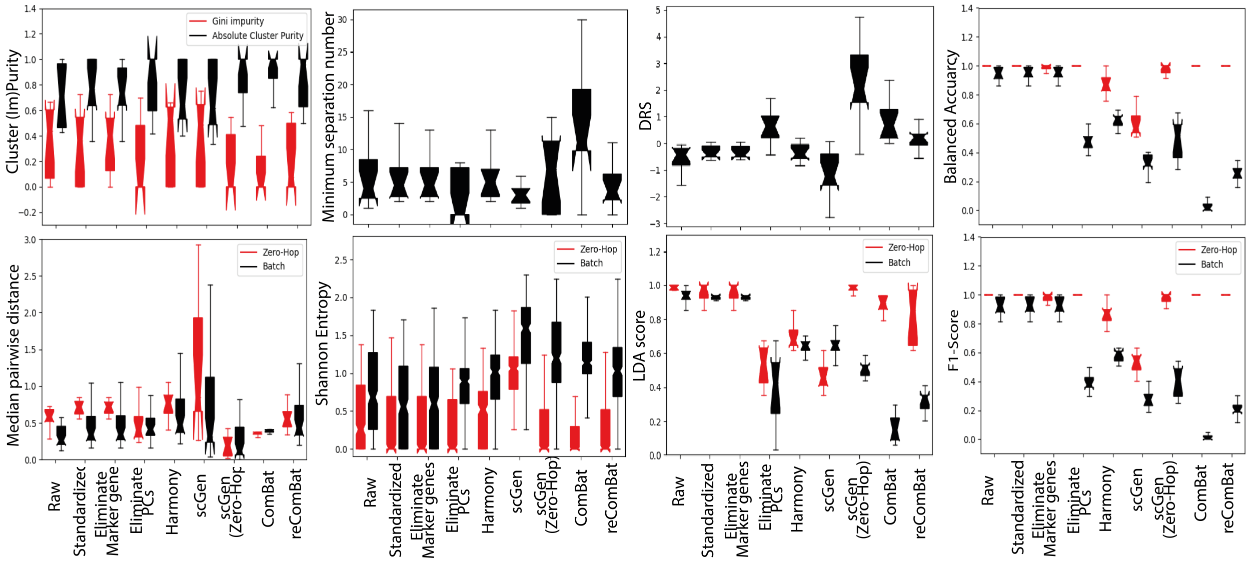


**Fig. S17.** Evaluation metrics scoring the impact of batch-effects by evaluating the variety of different batches and/or Zero-Hops of the (un-) corrected bulkRNAsq data. Box plots represent the lower and upper quartiles (box) together with the median (central dents) and full range (whiskers) over all samples, clusters, or Zero-Hops depending on the relevant metric. LDA scores and LR classification performance are reported over ten cross validation folds.

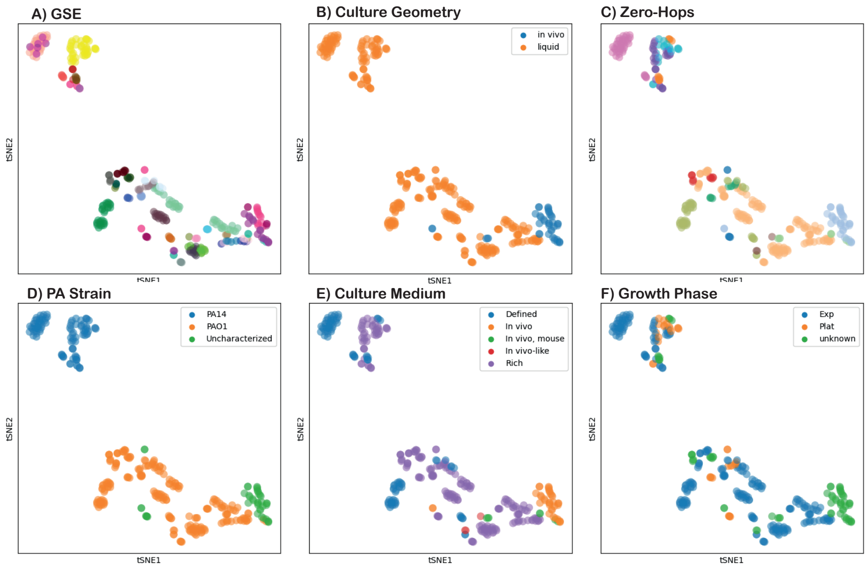


**Fig. S18.** tSNE plots of the uncorrected bulkRNAsq data colored by batches (A), all evaluated experimental designs (B-H), and the defined Zero-Hops (I). Clustering is largely driven by the GSE (i.e. batch) rather than by the underlying culture conditions or microbial strains.

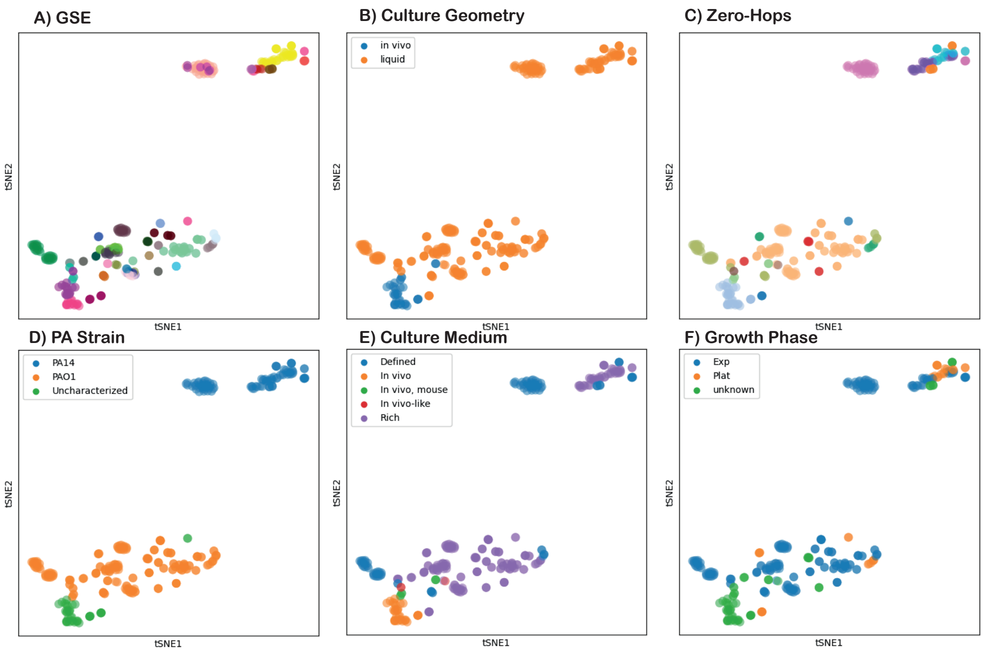


**Fig. S19.** tSNE plots of the corrected bulkRNAsq data using Z-scoring (i.e. standardization) colored by batches (i.e. GSE) (A), all evaluated experimental designs (B-H), and the defined Zero-Hops (I).

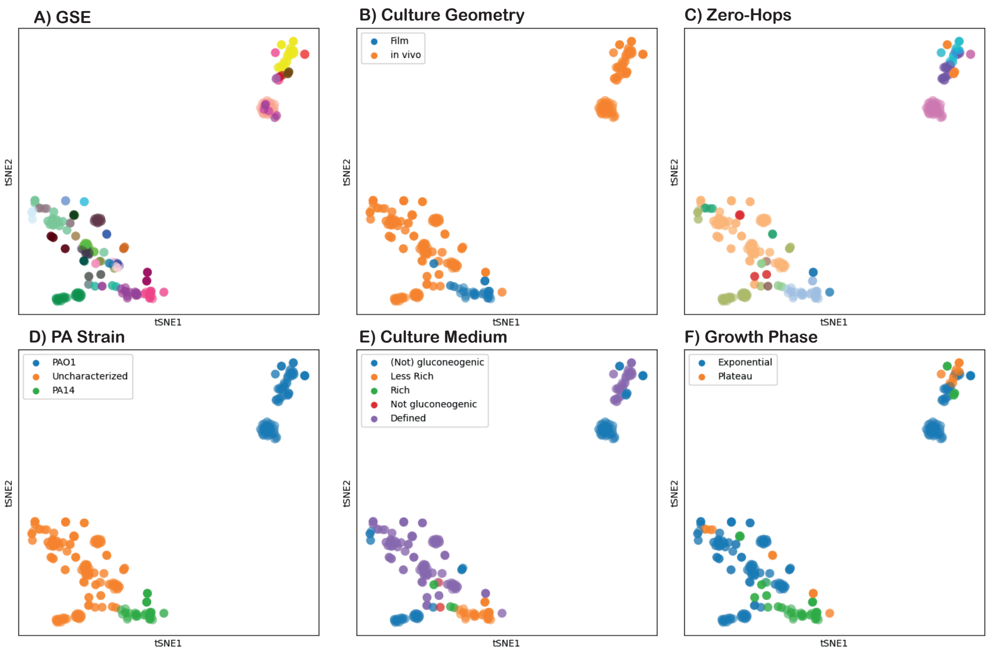


**Fig. S20.** tSNE plots of the corrected bulkRNAsq data using marker gene elimination colored by batches (i.e. GSE)(A), all evaluated experimental designs (B-H), and the defined Zero-Hops (I).

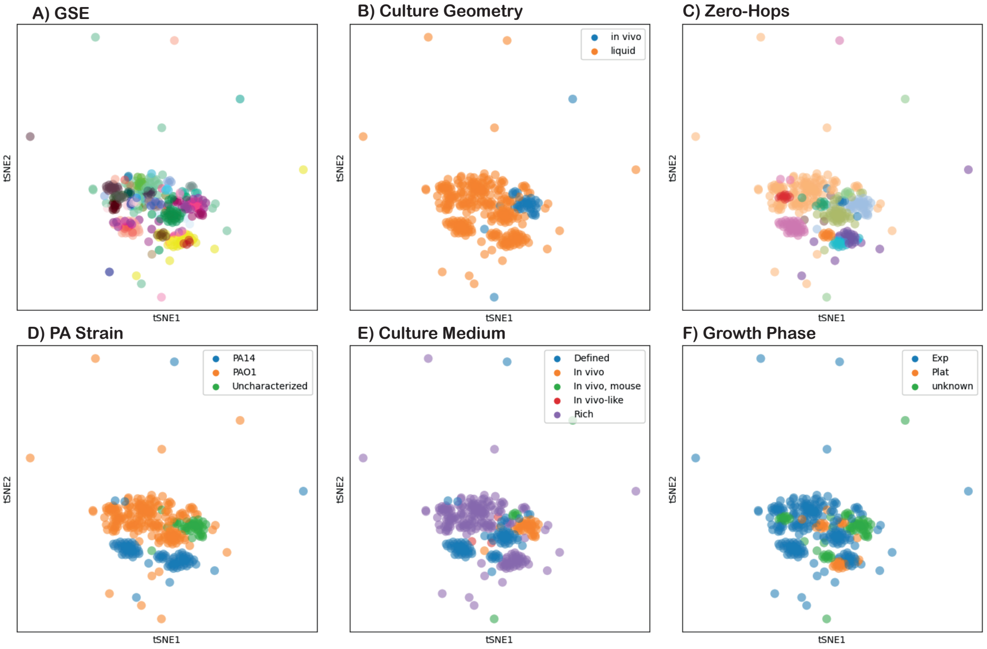


**Fig. S21.** tSNE plots of the corrected bulkRNAsq data using PC elimination coloured by batches (i.e. GSE)(A), all evaluated experimental designs (B-H), and the defined Zero-Hops (I).

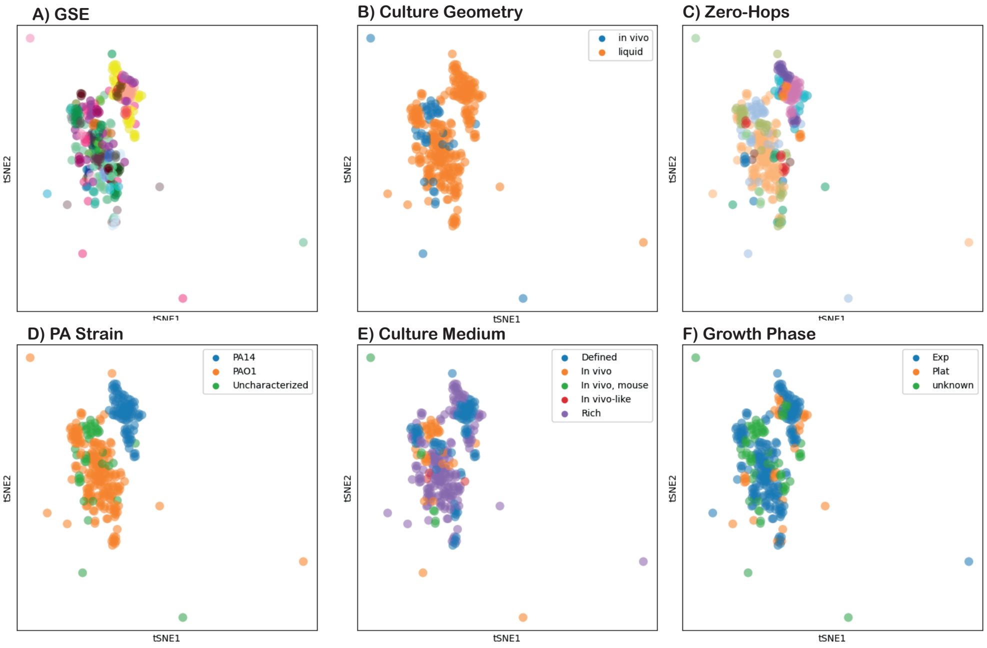


**Fig. S22.** tSNE plots of the corrected bulkRNAsq data using Harmony colored by batches (i.e. GSE)(A), all evaluated experimental designs (B-H), and the defined Zero-Hops (I).

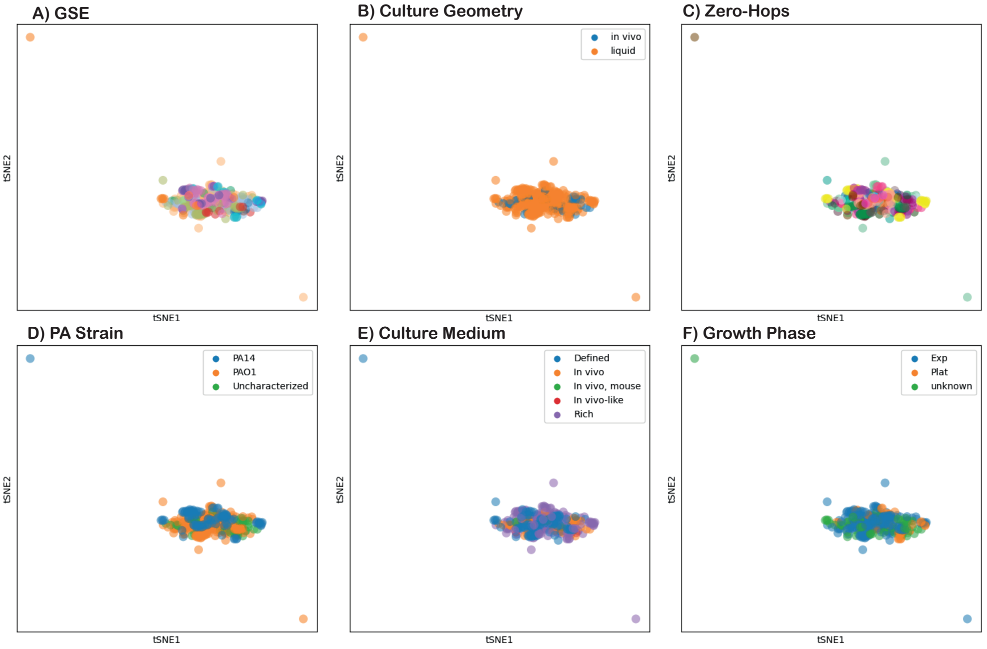


**Fig. S23.** tSNE plots of the corrected bulkRNAsq data using scGen without information on Zero-Hops. Plots are colored by batches (i.e. GSE)(A), all evaluated experimental designs (B-H), and the defined Zero-Hops (I).

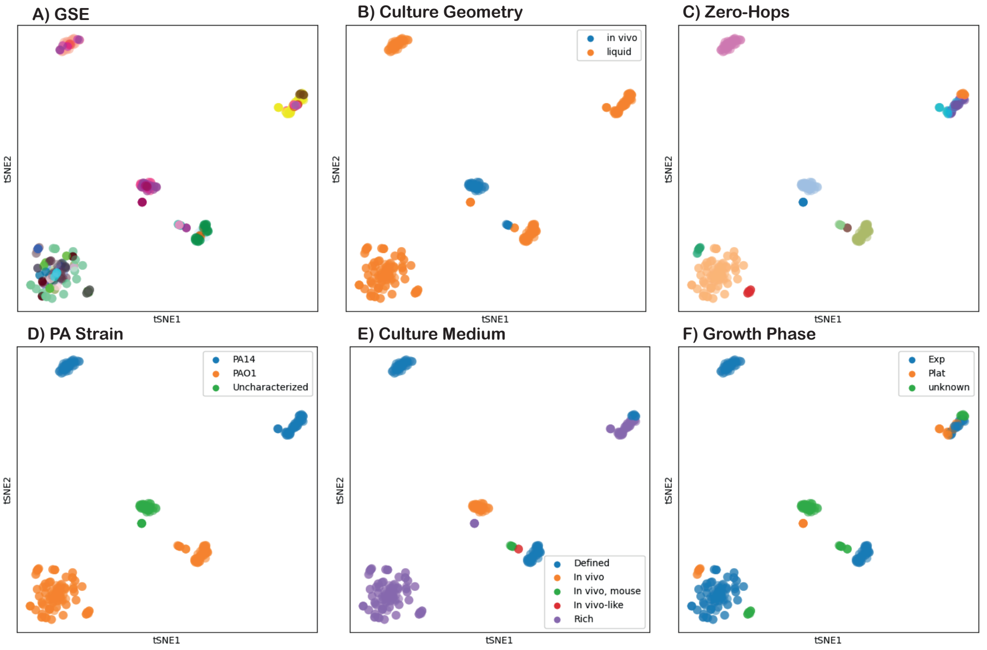


**Fig. S24.** tSNE plots of the corrected bulkRNAsq data using scGen with Zero-Hop used as label. Plots are colored by batches (i.e. GSE)(A), all evaluated experimental designs (B-H), and the defined Zero-Hops (I).

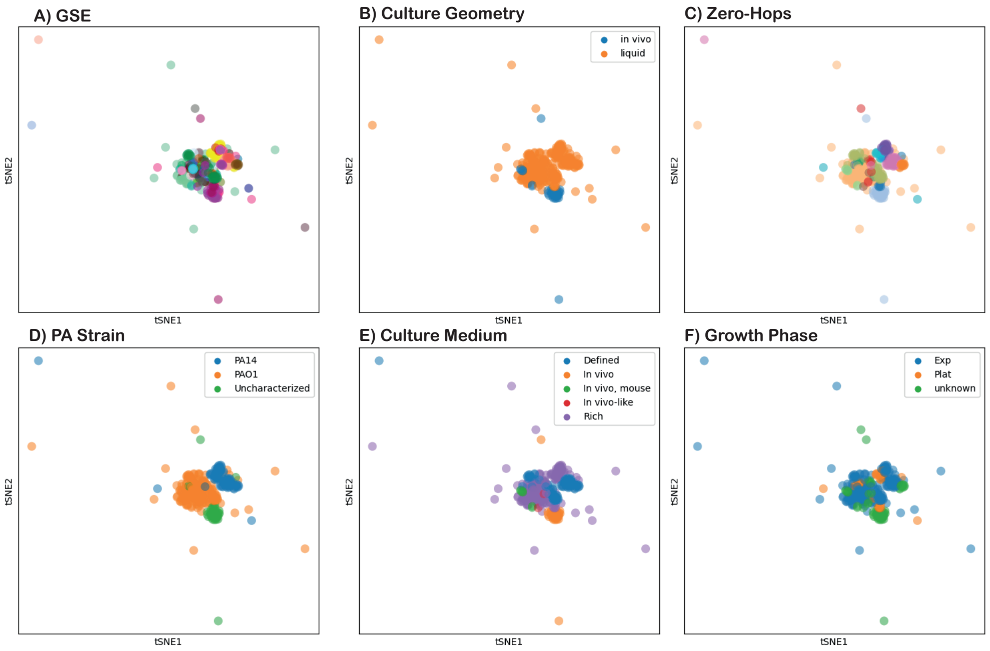


**Fig. S25.** tSNE plots of the corrected bulkRNAsq data using ComBat. This correction did not yield a unique solution. Plots are colored by batches (i.e. GSE)(A), all evaluated experimental designs (B-H), and the defined Zero-Hops (I).

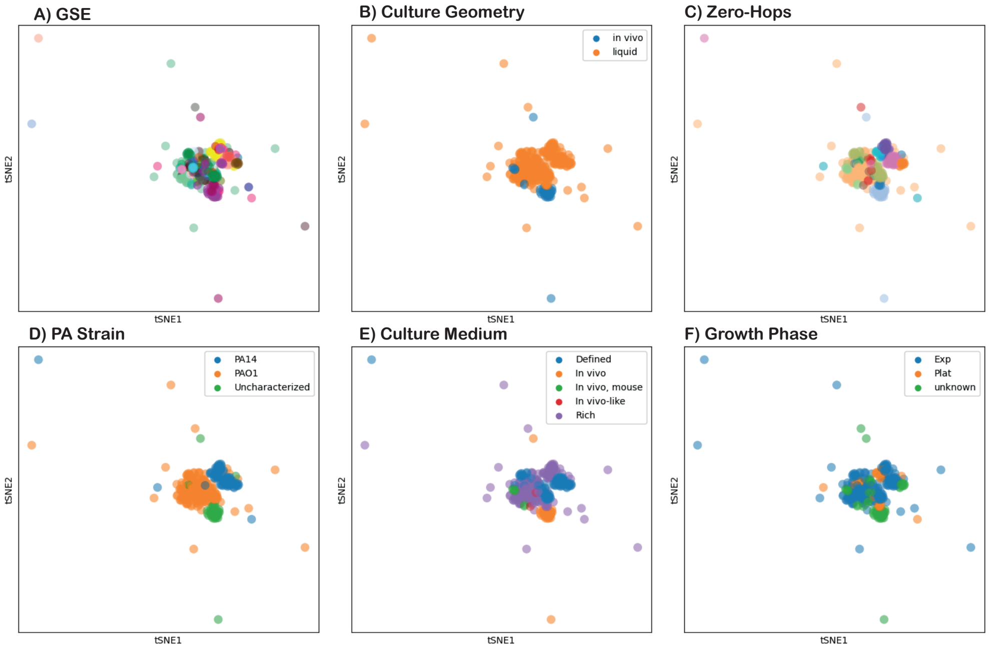


**Fig. S26.** tSNE plots of the corrected bulkRNAsq data using ridge *reComBat* with a regularisation strength $\lambda_2 = 1e-9$ colored by batches (i.e. GSE) (A), all evaluated experimental designs (B-H), and the defined Zero-Hops (I).

### G Evaluation of differences in gene expression between Zero-Hops in micro array data

Figure S27 gives an overview of the top 50 ranked genes driving differences in expression profiles between representative examples of Zero-Hops.

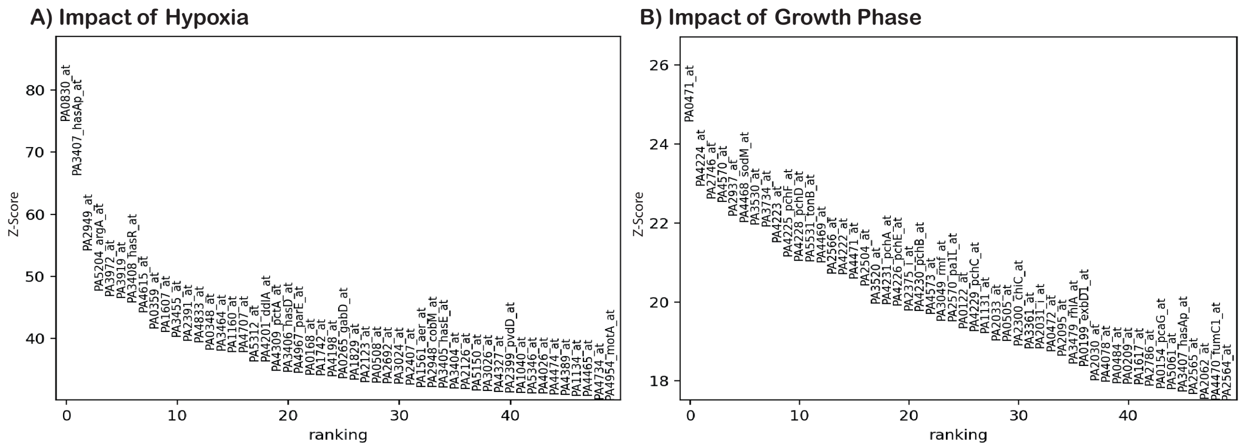


| Zero-Hop | Strain | Phase | Culture | T [°C] | Oxygenation | Medium | Antibiotic | $n_{Samples}$ | $n_{GSE}$ |
|---|---|---|---|---|---|---|---|---|---|
| 0 | PAO1 | Exp. | liquid | 35-38 | hypoxic | rich | no | 48 | 2 |
| 35 | PAO1 | Exp. | liquid | 35-38 | aerobic | rich | no | 170 | 23 |

| Zero-Hop | Strain | Phase | Culture | T [°C] | Oxygenation | Medium | Antibiotic | $n_{Samples}$ | $n_{GSE}$ |
|---|---|---|---|---|---|---|---|---|---|
| 33 | PAO1 | Plateau | liquid | 35-38 | aerobic | rich | no | 38 | 9 |
| 35 | PAO1 | Exp. | liquid | 35-38 | aerobic | rich | no | 170 | 23 |

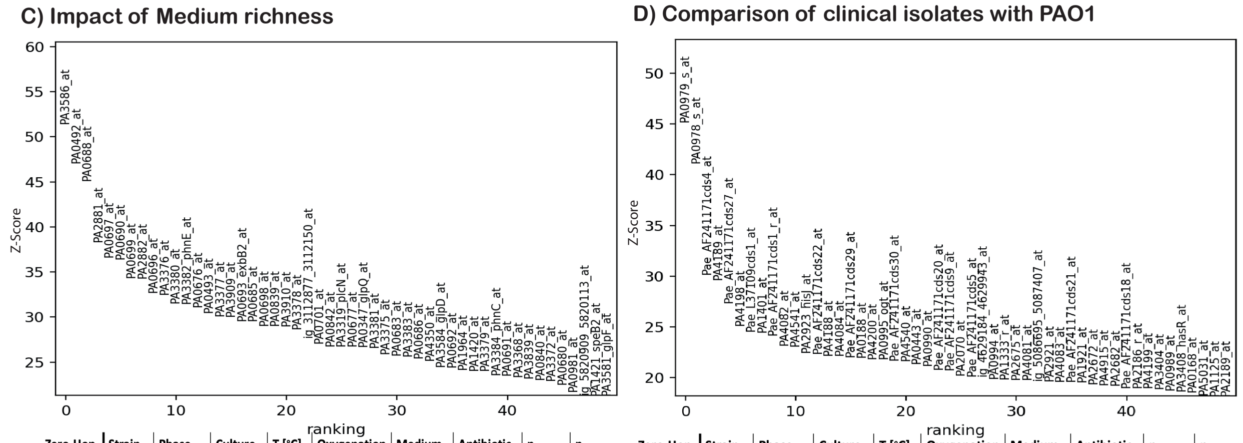


| Zero-Hop | Strain | Phase | Culture | T [°C] | Oxygenation | Medium | Antibiotic | $n_{Samples}$ | $n_{GSE}$ |
|---|---|---|---|---|---|---|---|---|---|
| 11 | PAO1 | Exp. | liquid | 35-38 | aerobic | less rich | no | 48 | 2 |
| 35 | PAO1 | Exp. | liquid | 35-38 | aerobic | rich | no | 170 | 23 |

| Zero-Hop | Strain | Phase | Culture | T [°C] | Oxygenation | Medium | Antibiotic | $n_{Samples}$ | $n_{GSE}$ |
|---|---|---|---|---|---|---|---|---|---|
| 16 | Clinical | Exp. | liquid | 35-38 | aerobic | rich | no | 114 | 7 |
| 35 | PAO1 | Exp. | liquid | 35-38 | aerobic | rich | no | 170 | 23 |

**Fig. S27.** Visualisation of the ranking of the top 50 genes differing in their expression between the indicated Zero-Hops. Key pathways affected by variations in culture condition or antibiotic treatment are greatly associated with previously reported findings indicating that biologically relevant variation was retained by *reComBat* ($\lambda_2$ = 1e-9,$\lambda_1$ = 0).

### References

1. Afgan, E., Baker, D., Batut, B., van den Beek, M., Bouvier, D., Čech, M., Chilton, J., Clements, D., Coraor, N., Grüning, B.A., Guerler, A., Hillman-Jackson, J., Hiltemann, S., Jalili, V., Rasche, H., Soranzo, N., Goecks, J., Taylor, J., Nekrutenko, A., Blankenberg, D.: The Galaxy platform for accessible, reproducible

and collaborative biomedical analyses: 2018 update. Nucleic Acids Research **46**(W1), W537–W544 (jul 2018). https://doi.org/10.1093/nar/gky379
2. Anders, S., Pyl, P.T., Huber, W.: HTSeq–a Python framework to work with high-throughput sequencing data. Bioinformatics **31**(2), 166–169 (jan 2015). https://doi.org/10.1093/bioinformatics/btu638
3. Barrett, T., Wilhite, S.E., Ledoux, P., Evangelista, C., Kim, I.F., Tomashevsky, M., Marshall, K.A., Phillippy, K.H., Sherman, P.M., Holko, M., Yefanov, A., Lee, H., Zhang, N., Robertson, C.L., Serova, N., Davis, S., Soboleva, A.: NCBI GEO: archive for functional genomics data sets–update. Nucleic acids research **41**(Database issue), D991–5 (jan 2013). https://doi.org/10.1093/nar/gks1193
4. Bolger, A.M., Lohse, M., Usadel, B.: Trimmomatic: a flexible trimmer for Illumina sequence data. Bioinformatics **30**(15), 2114–2120 (aug 2014). https://doi.org/10.1093/bioinformatics/btu170
5. Chazarra-Gil, R., van Dongen, S., Kiselev, V.Y., Hemberg, M.: Flexible comparison of batch correction methods for single-cell rna-seq using batchbench. Nucleic acids research **49**(7), e42–e42 (2021)
6. Hastie, T., Tibshirani, R., Friedman, J.: The Elements of Statistical Learning. Springer Series in Statistics, Springer New York Inc., New York, NY, USA (2001)
7. Johnson, W.E., Li, C., Rabinovic, A.: Adjusting batch effects in microarray expression data using empirical Bayes methods. Biostatistics **8**(1), 118–127 (04 2006). https://doi.org/10.1093/biostatistics/kxj037
8. Kim, D., Langmead, B., Salzberg, S.L.: HISAT: a fast spliced aligner with low memory requirements. Nature Methods **12**(4), 357–360 (apr 2015). https://doi.org/10.1038/nmeth.3317
9. Korsunsky, I., Millard, N., Fan, J., Slowikowski, K., Zhang, F., Wei, K., Baglaenko, Y., Brenner, M., Loh, P.r., Raychaudhuri, S.: Fast, sensitive and accurate integration of single-cell data with Harmony. Nature Methods **16**(12), 1289–1296 (dec 2019). https://doi.org/10.1038/s41592-019-0619-0
10. Liu, W., Li, Z.: An efficient parallel algorithm of n-hop neighborhoods on graphs in distributed environment. Frontiers of Computer Science **13**(6), 1309–1325 (2019)
11. Lotfollahi, M., Wolf, F.A., Theis, F.J.: scGen predicts single-cell perturbation responses. Nature methods **16**(8), 715–721 (2019). https://doi.org/10.1038/s41592-019-0494-8
12. Rong, Z., Tan, Q., Cao, L., Zhang, L., Deng, K., Huang, Y., Zhu, Z.J., Li, Z., Li, K.: NormAE: Deep Adversarial Learning Model to Remove Batch Effects in Liquid Chromatography Mass Spectrometry-Based Metabolomics Data. Analytical chemistry **92**(7), 5082–5090 (2020). https://doi.org/10.1021/acs.analchem.9b05460
13. Tran, H.T.N., Ang, K.S., Chevrier, M., Zhang, X., Lee, N.Y.S., Goh, M., Chen, J.: A benchmark of batch-effect correction methods for single-cell RNA sequencing data. Genome Biology **21**(1), 12 (dec 2020). https://doi.org/10.1186/s13059-019-1850-9
14. Varma, S.: Blind estimation and correction of microarray batch effect. PLOS ONE **15**(4), e0231446 (apr 2020). https://doi.org/10.1371/journal.pone.0231446
15. Whiteside, M.D., Winsor, G.L., Laird, M.R., Brinkman, F.S.L.: OrthologeDB: a bacterial and archaeal orthology resource for improved comparative genomic analysis. Nucleic Acids Research **41**(D1), D366–D376 (jan 2013). https://doi.org/10.1093/nar/gks1241
